# Depth and evenness of sequence coverage are associated with assembly quality, genome structure, and choice of sequencing platform in archived plastid genomes

**DOI:** 10.1101/2022.05.06.490930

**Authors:** Nils Jenke, Michael Gruenstaeudl

## Abstract

In plastid genomes, the depth and evenness of sequence coverage are considered important indicators for assembly quality. However, the precise manifestations that sequencing depth and evenness can have in the assembly of these genomes, as well as any differences across individual genome sections, have yet to be evaluated. This investigation aims to identify the impact that sequencing depth and evenness can have on the assembly of plastid genomes and how both metrics are related to plastid genome structure. Specifically, we assess if sequencing evenness and reduced sequencing depth have significant correlations with, or significant differences among, individual genome sections, assembly quality metrics, the sequencing platforms employed, and the software tools used for genome assembly. To that end, we retrieve published plastid genomes as well as their sequence reads and genome metadata from public databases, measure sequencing depth and evenness across their sequences, and test several hypotheses on genome assembly and structure through non-parametric statistical tests. The results of our analyses show significant differences in sequencing depth across the four structural partitions as well as between the coding and non-coding sections of the plastid genomes, a significant correlation between sequencing evenness and the number of ambiguous nucleotides per genome, and significant differences in sequencing evenness between various sequencing platforms. Based on these results, we conclude that the observed differences and correlations are not a product of chance alone but possibly genuine manifestations of sequencing depth and evenness during the assembly of these genomes.

## INTRODUCTION

The majority of publicly available plastid genomes have been generated through a multitude of different sequencing techniques and assembly methods, with each exhibiting an idiosyncratic error rate. More than ten thousand unique and complete plastid genomes are now available on NCBI Nucleotide (last accessed on 01-Mar-2021; Mehl and Gruenstaeudl 2020). Many of these genomes have been generated through Illumina DNA sequencing but other next-generation sequencing (NGS) platforms have been employed as well (Slatko et al., 2018). Most NGS platforms exhibit intrinsic sequencing error rates that range between 0.1% and 1% of generated nucleotides Fuller et al. (2009); Fox et al. (2014). A non-trivial number of nucleotides may, thus, be erroneous in a typical plastid genome assembly by default, even if not all sequencing errors become manifest in the final sequence. Researchers need to take such natural error rates into consideration, particularly when conducting genomic comparisons between closely related taxa. Similarly, a plethora of different software tools and bioinformatic pipelines for the *de novo* assembly of plastid genomes exist (e.g., Dierckxsens et al., 2016; Coissac et al., 2016; Ankenbrand et al., 2018; Jin et al., 2020). Many of these tools exhibit disparate abilities to assemble plastid genomes in full, and receiving complete, circular, and error-free sequences upon automated plastid genome assembly is still the exception rather than the rule (Alhakami et al., 2017; Freudenthal et al., 2020). In fact, most tools cannot assemble a complete plastid genome in one sitting but generate large sequence contigs that require post-processing to obtain a complete assembly (Jin et al., 2020). The IRs of the plastid genome, for example, are notorious in their interference with the assembly process (e.g., Logacheva et al., 2014) and often mandate the identification and correction of the precise IR boundaries as well as the manual concatenation of the contigs (Wu et al., 2015; Amiryousefi et al., 2018). Most assembly software is, thus, employed as part of bioinformatic workflows that combine the functionality of various software tools (Gruenstaeudl et al., 2018). However, the post-processing and scaffolding of contigs can introduce gaps and ambiguous nucleotides into the assembly products (Paulino et al., 2015) and, thus, increase the rate of assembly errors in the final genomes.

A considerable number of publicly available plastid genomes appear to exhibit suboptimal genome assemblies, highlighting the need for the increased application of assembly quality metrics. The dark figure of suboptimally assembled plastid genomes appears to be high. Wu et al. (2015), for example, reported that 51 out of the 424 plastid genomes then available on NCBI Nucleotide contained ambiguities. Similarly, (Mehl and Gruenstaeudl, 2020) reported that more than half of all plastid genomes on NCBI Nucleotide lacked annotations for their IR regions. Moreover, a considerable number of published plastid genomes contain non-identical IRs (Zheng et al., 2020, pers. obs.). Unfortunately, many of these suboptimal genome records remain unrecognized, as the assessment of assembly quality indicators is not mandated at or before genome publication. Several such indicators are readily available to researchers, and three types can be applied with ease to assess the assembly quality of plastid genomes. First, researchers can apply sequence contiguity metrics, which typically indicate the length of the shortest contig needed to cover a certain percentage (e.g., 50%) of the target sequence (Earl et al., 2001). These metrics are commonplace in the sequencing of nuclear genomes and are increasingly employed on plastid genomes (e.g., Gruenstaeudl et al., 2018; Zhu et al., 2018). Second, researchers can use the structure and gene synteny of plastid genomes as a proxy for their assembly quality given the high level of structural conservation among plastid genomes, at least among land plants (Mower and Vickrey, 2018). For example, equality in length and sequence of the IRs of a plastid genome can be used as an indicator of its overall assembly quality (Gruenstaeudl and Jenke, 2020; Zheng et al., 2020), as IR equality is maintained naturally and rarely deviated from (Goulding et al., 1996; Ruhlman et al., 2017). Third, researchers can measure the depth of sequence coverage of a genome assembly (Sims et al., 2014), with both the average sequencing depth and the evenness of coverage indicative of assembly quality (Chen et al., 2013; Wang et al., 2021). Average sequencing depth can assist in identifying the overall adequacy of sequence coverage, whereas the precise distribution of sequencing depth can help to identify regions of inadequate coverage. Among these quality indicators, the average genome-wide sequencing depth is employed most often in contemporary plastid genome studies (Twyford and Ness, 2017; McKain et al., 2018). For the remainder of this investigation, we will follow the terminology of Sims et al. (2014) and use the terms ‘sequencing depth’ and ‘sequence coverage’ interchangeably but primarily employ to former if the precise level of the latter is to be emphasized.

## INTRODUCTION

Suboptimal genome assemblies often exhibit uneven sequence coverage, which can be measured through different metrics. Reliable genome assemblies typically exhibit a relatively uniform sequence coverage, as all regions of the genome are covered by a similar amount of aligned sequence reads (Sims et al., 2014). Many *de novo* assembly algorithms even require a relatively even sequence read distribution for successful genome assembly (Peng et al., 2012; McCorrison et al., 2014). By contrast, many suboptimal genome assemblies exhibit an uneven sequence coverage, as mis-assembled regions are poorly, if at all, covered by sequence reads (Peng et al., 2012). Numerous reports on this phenomenon exist. (Ekblom et al., 2014), for example, reported a negative correlation between erroneous base calls and sequencing evenness in the assembly of vertebrate mitochondria. They also found that genomic regions with a high GC content predominantly exhibited uneven sequence coverage, corroborating an earlier report of Benjamini and Speed (2012). Similarly, Sassenhagen and Rengefors (2019) identified inconsistencies in the plastid genome assemblies of heterokont algae and linked them to uneven sequence coverage. Moreover, Scheunert et al. (2020) concluded the need for caution regarding uneven sequence coverage in a study on long-read sequencing and assembly of sunflower plastid genomes. Evenness in sequence coverage (hereafter ‘sequencing evenness’) should, thus, be among the standard indicators for assessing the assembly quality of plastid genomes (Mokry et al., 2010; Gruenstaeudl and Jenke, 2020). Gu et al. (2016), for example, employed sequencing evenness as an indicator to refine the assembly of the plastid genome of crape myrtle (*Lagerstroemia fauriei*). Similarly, Wang et al. (2021) assessed sequencing evenness in the nuclear genome assembly of a tomato cultivar to infer the factors contributing to genome mis-assembly. Several metrics to measure sequencing evenness exist, with the standard deviation (*σ*) around the mean value of the normalized sequence coverage representing a commonly used metric (Oexle, 2016). Users may alternatively employ the evenness score (‘E-score’; Mokry et al., 2010), which measures sequencing evenness over a target sequence, was specifically designed for quick computational inference (Oexle, 2016), and represents the primary method to assess sequencing evenness in this investigation. Despite the simplicity to measure sequencing evenness, few investigations have employed this measure in studies on plastid genomes. The precise effect of sequencing evenness on the assembly of plastid genomes, thus, remains poorly understood.

The present investigation aims to identify the impact that sequencing depth and sequencing evenness can have on the assembly of plastid genomes and how both metrics are related to the structure of these genomes. Specifically, we aim to explore if sequencing evenness and reduced sequencing depth have significant correlations with, or significant differences among, factors that characterize plastid genome assembly and structure. To that end, we set up two hypotheses regarding sequencing depth (i.e., hypotheses a and b) and four hypotheses regarding sequencing evenness (i.e., hypotheses c–f). The hypotheses on sequencing depth aim to identify significant differences across structural genome partitions and between coding and non-coding genome sections. They are: (a) sequencing depth is not significantly different across the four structural partitions of the plastid genome (i.e., LSC, IR_B_, SSC, and IR_A_); and (b) sequencing depth is not significantly different between coding versus non-coding sections of the plastid genome. To focus on cases of reduced sequence coverage, sequencing depth is not employed directly but substituted with the number of genome regions with reduced sequence coverage in relation to the average genome-wide sequencing depth. The hypotheses on sequencing evenness aim to identify correlations with assembly quality metrics as well as significant differences across sequencing platforms and assembly software tools. They are: (c) sequencing evenness of a plastid genome is not correlated with the number of its ambiguous nucleotides; (d) sequencing evenness of a plastid genome is not correlated with the number of sequence mismatches between its IRs; (e) sequencing evenness is not significantly different across sequencing platforms; and (f) sequencing evenness is not significantly different across assembly software tools. To evaluate these hypotheses, we test them on a large set of archived plastid genomes. Specifically, we retrieve complete plastid genomes stored on NCBI Nucleotide for which raw sequence reads and various types of metadata are accessible, assess the data dispersion of sequencing depth and evenness across these genomes, and then test our hypotheses in a non-parametric statistical framework.

## MATERIALS AND METHODS

### Genome sequences and metadata

To test the hypotheses of this investigation, we selected 194 genomes of the plastid genome compilation of Freudenthal et al. (2020). Our selection was based on two conditions: (i) both sequencing depth and sequencing evenness needed to be measurable on the genomes, and (ii) the genomes needed to represent seed plants. The first condition reflected the requirements that the raw sequence reads of each genome were retrievable from NCBI SRA (Kodama et al., 2011), that each genome was parse-able via the R package genbankr (which represents the primary parsing tool for GenBank flatfiles in R; Becker and Lawrence, 2020), and that none of the genomes exhibited large gaps in their sequence coverage, which would preclude the calculation of sequencing evenness. The second condition reflected the requirement of a quadripartite genome structure with a similar gene content across the four genome partitions, which is a precondition to test hypothesis (a) and generally satisfied by the plastid genomes of seed plants (Mower and Vickrey, 2018). The 194 plastid genomes so selected represented various lineages of angio- and gymnosperms and the products of different researchers, sequencing technologies, and assembly methods. The accession numbers (NCBI Nucleotide and NCBI SRA) of each genome as well as the plant species it represents are given in Supplementary Table 1. To compile these genomes and their raw sequence reads into a single dataset, the complete sequence of each genome was retrieved from NCBI Nucleotide with Entrez Direct v.13.9 (Kans, 2013) and the corresponding sequence reads from NCBI SRA with SRA Toolkit v.2.10.8 (SRA Toolkit Development Team, 2020). To replicate the likely quality filtering conducted during the original assembly of the genomes, we evaluated the sequence quality and successful pairing of the reads with Trimmomatic v.0.39 (Bolger et al., 2014) and kept only paired reads longer than 36 bp after removal of all terminal nucleotides below a quality threshold of 3. In light of the observation that IR annotations of archived plastid genomes are not always correct (Mehl and Gruenstaeudl, 2020), we also evaluated and, where necessary, corrected the exact start and end positions of the IRs of the plastid genomes using script 4 of the workflow of Gruenstaeudl et al. (2018) but we did not infer new IR annotations where such were not originally listed. To test the hypotheses of this investigation, we also extracted two classes of metadata from the plastid genomes under study. For each genome, we extracted the name of the sequencing platform from NCBI SRA and the name of the assembly software employed from NCBI Nucleotide. The retrieval of this data required several cycles of data parsing, as the names and version numbers were not specified uniformly across records. Specifically, we had to remove spelling mistakes and homogenize the specification of version numbers to achieve comparability across genomes. All metadata values extracted are given in Supplementary Table 2.

### Sequencing depth and evenness

For each plastid genome under study, we calculated both sequencing depth and sequencing evenness. Sequencing depth was calculated through a two-step procedure. First, the quality-filtered sequence reads of each genome were mapped onto the complete genome sequence using Bowtie2 v.2.4.2 (Langmead and Salzberg, 2012) under default settings. The resulting read mapping was sorted and indexed with Samtools v.1.9 (Li et al., 2009) and saved as a BAM file. Second, the sequencing depth of each genome was calculated using PACVr v.1.0.1 (Gruenstaeudl and Jenke, 2020) by supplying the complete genome sequence plus sequence annotations as GenBank flatfile and the read mapping as corresponding BAM file. A default window size of 250 bp was employed. Coverage windows whose sequencing depth was one standard deviation below the average genome-wide sequencing depth were classified as ‘windows with reduced sequencing depth’ (abbreviated as ‘WRSD’) and categorized by their genome partition (i.e., LSC, IRB, SSC, or IR_A_) and coding status (i.e., coding versus non-coding DNA). Sequencing evenness was calculated as a genome-wide E-score value with PACVr based on the inferred sequencing depth profiles and using the equation of Oexle (2016). Under these conditions, the E-score ranges between 0 and 1, with values close to 1 indicating even coverage and values close to 0 indicating uneven coverage. Preliminary analyses indicated that E-score values equal to, or smaller than, 0.80 were already representative of highly uneven sequence coverage. E-score values that represent an uneven sequence coverage and are more than 1.5 × *interquartile* range below the first quartile value of the E-score distribution are considered outliers. The number of WRSD per genome partition and per coding status as well as the E-score value for each plastid genome are given in Supplementary Table 2.

### Assembly quality metrics

For each plastid genome under study, we calculated two assembly quality metrics: the number of ambiguous nucleotides per genome and the number of mismatches between the IRs. Both quality measures have been used as indicators of assembly accuracy and completeness (Liu et al., 2018; Earl et al., 2001; Shin and Park, 2014; Gruenstaeudl et al., 2017). The number of ambiguous nucleotides per genome sequence is a direct measure of assembly quality, with a lower number indicating a higher assembly quality. The number of mismatches between the two IR sequences of a plastid genome is an indirect measure of assembly quality, with a lower number indicating greater IR similarity and, by extension, a higher assembly quality. Both metrics were calculated with PACVr. Prior to calculating IR mismatches, we verified equality between the IRs in gene content and order; genomes that lacked gene synteny among their IRs were disqualified from further analysis. Upon calculation of both metrics, we standardized their values through a transformation following Tukey’s ladder of powers (Tukey, 1977); the resulting values ranged between 0 and 1, with values close to 1 indicating high assembly quality and values close to 0 indicating low assembly quality. This transformation would not affect the outcome of our statistical tests but simplified the comparability of their results. All inferred metric values prior to transformation are given in Supplementary Table 2.

### Statistical analysis

To test our hypotheses in a statistical framework, we classified the compiled data into five quantitative and two categorical variables. The quantitative variables comprised the genome-wide E-score value, the number of ambiguous nucleotides per genome, the number of IR sequence mismatches, and the number of WRSD per genome partition and per coding status. The categorical variables comprised the sequencing platform and the assembly software tool employed. To ensure a balanced calculation of the test statistics among the categorical variables (Ostertagova et al., 2014), we defined a minimum sample size of five genome records for each state; any state that did not meet this sample size was removed and its members subsumed into the state ‘missing’. The assumption of homoscedasticity was evaluated in all variables through Levene’s test (Levene, 1960). Homoscedasticity was confirmed for all variables that represented, or were compared against, sequencing evenness but not for variables that were compared against sequencing depth; our selection of statistical tests was designed accordingly.

The hypotheses of this investigation were tested with five different statistical tests. Hypotheses (a), (e), and (f) were evaluated using a non-parametric one-way analysis of variance to identify differences between three or more independent groups. Specifically, we employed the Kruskal-Wallis test (Kruskal and Wallis, 1952) to assess if the distributions of the number of WRSD were significantly different among the four genome partitions, if the distributions of E-score values were significantly different among different sequencing platforms, and if the distribution of E-score values were significantly different among different assembly software. Significant differences between the genome partitions, sequencing platforms, and assembly software were also evaluated through pairwise comparisons using post-hoc pairwise Wilcoxon rank-sum tests (Armstrong and Hilton, 2010). To that end, Benjamini-Hochberg corrections (Benjamini and Hochberg, 1995) were applied on the test data to mitigate the problem of multiple testing (Sainani, 2009). Hypothesis (b) was also evaluated using a non-parametric test to compare differences between two independent groups. Specifically, we employed the Wilcoxon rank-sum test (Wilcoxon, 1992) to assess if the distributions of the number of WRSD of the coding and non-coding genome sections were significantly different. Due to the presence of ties in the data, the test statistic values were inferred through normal approximation. To evaluate the relevance of the p-values generated through the Kruskal-Wallis and the Wilcoxon rank-sum tests (LeCroy and Krysik, 2007), we additionally assessed the statistical effect size using Cohens-d (Cohen, 1977) for all comparisons conducted when testing hypotheses (a), (b), (e), and (f). P-values less than *α* = 0.05 were considered as significant, p-values less than *α* = 0.01 as very significant, and p-values less than *α* = 0.001 as highly significant in rejecting the null hypothesis. Effect size *d_c_* was considered to be small if *d_c_* ≥ 0.2, moderate if *d_c_* ≥ 0.5, and large if *d_c_* ≥ 0.8. Hypotheses (c) and (d) were evaluated using non-linear correlation tests. Specifically, we tested whether the E-score was correlated with either the number of ambiguous nucleotides or the number of IR mismatches at a significant level using Spearman’s rank statistic (*ρ*; Spearman, 2010). The correlation coefficient *R_s_* was considered to be small if *R_s_* ≥ 0.1, moderate if *R_s_* ≥ 0.3, and large if *R_s_* ≥ 0.5.

## RESULTS

### Genome metadata

Considerable heterogeneity was observed among the metadata values extracted from the plastid genomes (Table 1). The name of the sequencing platform was successfully extracted for all but one plastid genome and aggregated into 12 distinct states. Of these, four did not meet the sample size requirement, resulting in eight final states with a cumulative sample size of 187 genomes (96.5%). The sequencing platforms Illumina HiSeq 2000, Illumina HiSeq 2500, and Illumina MiSeq were observed with a high frequency. The name of the assembly software employed was successfully extracted for 74 of the plastid genomes and aggregated in 15 distinct states. Of these, ten did not meet the sample size requirement, resulting in five final states with a cumulative sample size of 54 genomes (27.8%). The assembly software tools Velvet (Zerbino and Birney, 2008) and CLC Genomics Workbench (CLC Bio, 2020) were observed with a high frequency.

**Table 1.** Overview of the relative frequency of different variable states of sequencing platform and assembly software and their respective E-scores. Prefix ‘‘Illumina” was redundant and, thus, removed from the names of sequencing platforms. Abbreviations used: Asm. = Assembly; GA. = Genome Analyzer; Seq. = Sequencing.

| Variable | State | n | % | $\Sigma$ % | E-score | |
| --- | --- | --- | --- | --- | --- | --- |
| | | | | | $\bar{x}$ | s |
| Seq. platform | GA II | 6 | 3.1 | 3.1 | 0.89 | 0.05 |
|  | HiSeq 1500 | 6 | 3.1 | 6.2 | 0.95 | 0.02 |
|  | HiSeq 2000 | 45 | 23.2 | 29.4 | 0.90 | 0.14 |
|  | HiSeq 2500 | 45 | 23.2 | 52.6 | 0.92 | 0.12 |
|  | HiSeq 4000 | 19 | 9.8 | 62.4 | 0.91 | 0.03 |
|  | HiSeq X Ten | 5 | 2.6 | 65.0 | 0.90 | 0.06 |
|  | MiSeq | 55 | 28.4 | 93.4 | 0.89 | 0.14 |
|  | NextSeq 500 | 6 | 3.1 | 96.5 | 0.84 | 0.11 |
|  | missing | 7 | 3.6 | 100 | 0.83 | 0.09 |
|  | *all* | 194 | 100 |  |  |  |
| Asm. softw. | CLC Asm. | 16 | 8.2 | 8.2 | 0.89 | 0.19 |
|  | Geneious | 5 | 2.6 | 10.8 | 0.84 | 0.23 |
|  | SOAPdenovo | 8 | 4.1 | 14.9 | 0.89 | 0.11 |
|  | Velvet | 19 | 9.8 | 24.7 | 0.92 | 0.04 |
|  | YASRA | 6 | 3.1 | 27.8 | 0.89 | 0.07 |
|  | missing | 140 | 72.2 | 100 | 0.90 | 0.12 |
|  | *all* | 194 | 100 |  |  |  |

### Sequencing depth

Considerable differences between the distributions of the number of WRSD were observed across the four genome partitions and across the coding and non-coding sections of the plastid genomes (Table 2). WRSD were identified across all four genome partitions in 165 (85.1%) of the plastid genomes and exhibited a mean of 46.99 across all LSC, 15.34 across all IR_B_, 9.66 across all SSC, and 12.64 across all IR_A_. The median value was higher than the mean value for each partition, indicating a negative skew in the WRSD distributions. A graphical comparison of the data dispersion of the number of WRSD across the four partitions indicated that the WRSD in the LSC were generally more numerous than in the other partitions (Figure 1A). For the coding and non-coding genome sections, WRSD were inferred in all plastid genomes and exhibited a mean of 44.66 across all coding and 16.90 across all non-coding sections. The comparison of the median and mean values indicated a negative skew in the WRSD distribution of the coding and a positive skew in the WRSD distribution of the non-coding sections of the genomes. A graphical comparison of the data dispersion in these two distributions indicated that the WRSD in the coding sections were generally more numerous than in the non-coding sections (Figure 1B).

**Figure 1.**
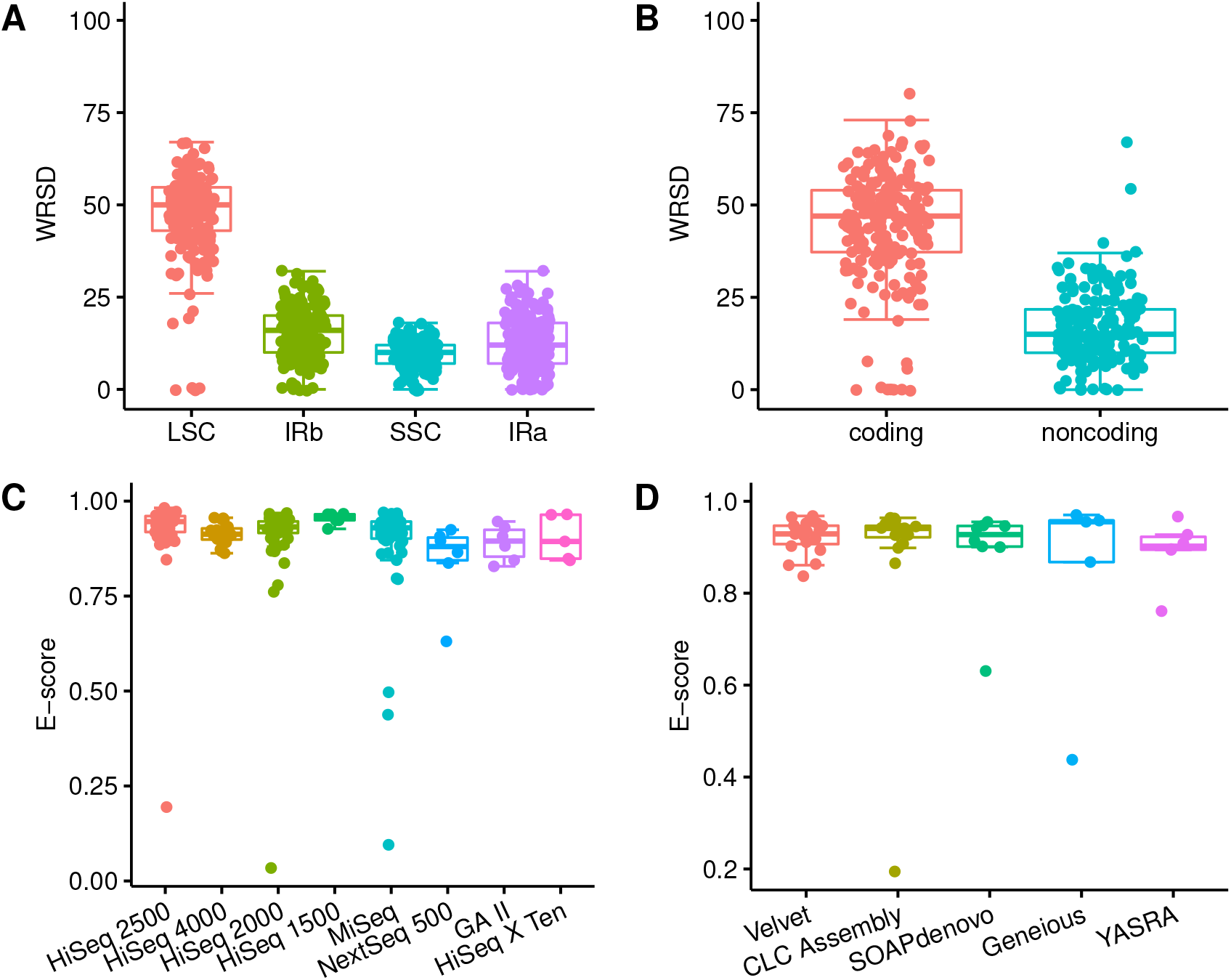
Visualization of the data dispersion of the number of WRSD among (A) the four genome partitions and (B) the coding and non-coding sections of the plastid genomes, as well as of the E-score values in relation to (C) different sequencing platforms and (D) different assembly software. Different states are color-coded for easier visualization.

**Table 2.** Overview of the data dispersion of the number of WRSD in relation to genome partitions and coding sections, of the E-scores, and of the two metrics on assembly quality. Column ‘Na’ lists the number of genomes in which the observed genome partition was not annotated and the inference of WRSD or IR mismatches, thus, not possible. Abbreviations used: Cod. = Coding; Nucl. = Nucleotides. All other abbreviations are as in Table 1.

| | Variable | n | Na | Min | q<sub>1</sub> | Median | $\bar{x}$ | q<sub>3</sub> | Max | s | IQR |
| --- | --- | --- | --- | --- | --- | --- | --- | --- | --- | --- | --- |
| Seq. depth | WRSD LSC | 166 | 28 | 0 | 43.00 | 50 | 46.99 | 54.75 | 67 | 12.22 | 11.75 |
|  | WRSD IR<sub>B</sub> | 167 | 27 | 0 | 10.00 | 16 | 15.34 | 20.00 | 32 | 7.13 | 10.00 |
|  | WRSD SSC | 166 | 28 | 0 | 7.00 | 10 | 9.66 | 12.00 | 18 | 3.71 | 5.00 |
|  | WRSD IR<sub>A</sub> | 165 | 29 | 0 | 7.00 | 12 | 12.64 | 18.00 | 32 | 7.04 | 11.00 |
|  | WRSD Cod. | 194 | 0 | 0 | 37.25 | 47 | 44.66 | 54.00 | 80 | 14.80 | 16.75 |
|  | WRSD Non-cod. | 194 | 0 | 0 | 10.00 | 15 | 16.90 | 21.75 | 67 | 9.67 | 11.75 |
| Seq. evenness | E-score | 194 | 0 | 0.03 | 0.90 | 0.93 | 0.90 | 0.95 | 0.98 | 0.12 | 0.05 |
| Asm. quality | Ambiguous nucl. | 194 | 0 | 0 | 0.00 | 0 | 6.43 | 0.00 | 551 | 44.63 | 0.00 |
|  | IR mismatches | 165 | 29 | 0 | 0.00 | 0 | 79.34 | 0.00 | 12,830 | 998.71 | 0.00 |

### Sequencing evenness

Considerable differences between the genome-wide E-score values of different plastid genomes were observed, with some genomes representing outliers with regard to the overall E-score data dispersion. The genome-wide E-score was inferred for all plastid genomes under study and exhibited a mean value of 0.90, with the center half of all values ranging between 0.90 to 0.95 (interquartile range = 0.05; Table 2). The median value was hereby higher than the mean value, indicating a negative skew in the E-score distribution. The highest E-score was detected in the plastid genome of *Asparagus officinalis* (accession NC_034777; E-score = 0.98), the smallest in the plastid genome of *Elaeis guineensis* (NC_017602; E-score = 0.03). The latter genome was one of 13 plastid genomes that were identified as outliers with regard to the E-score data dispersion across all genomes (Figure 2A; outliers labeled in red). These outlier genomes exhibited E-scores equal to, or less than, 0.80, which indicates a highly uneven sequence coverage. To illustrate the degree of uneven sequence coverage in these outliers, their genome-wide sequence coverage is visualized in Figure 2B and Supplementary Figures 1–13.

**Figure 2.**
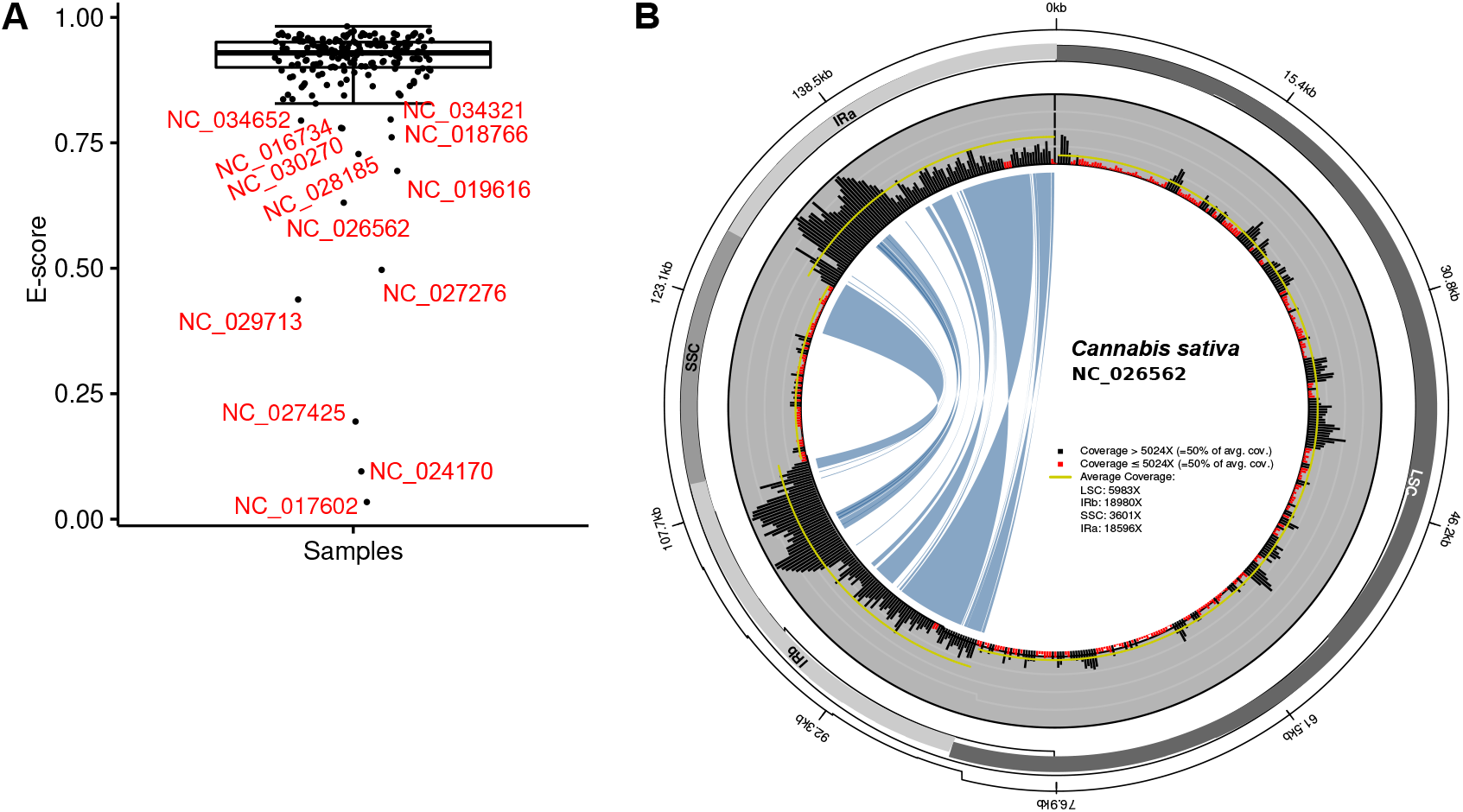
Data dispersion of E-score values and an example of a plastid genome with highly uneven sequence coverage. (A) Visualization of the data dispersion of the E-score values of all plastid genomes under study. The E-scores of genomes identified as outliers are labeled with their accession numbers in red. (B) Genome-wide visualization of the highly uneven sequence coverage of the plastid genome of *Cannabis sativa* (accession NC_026562). This genome was classified as an outlier with regard to its E-score value. Coverage calculation windows with less than half of the mean genome-wide sequencing depth are highlighted by red histogram bars.

### Assembly quality metrics

Large differences among the values of the two assembly quality metrics were observed across the plastid genomes (Table 2). The number of ambiguous nucleotides per genome ranged from none to 551 nucleotides, with the latter value found in the plastid genome of *Stachys byzantina* (accession NC_029825; Supplementary Table 2). The number of sequence mismatches between the two IRs of a plastid genome ranged from none to a total of 12,830, with the latter value found in the plastid genome of *Dendrobium nobile* (NC_029456). This extreme number of sequence mismatches between the IRs of the plastid genome record of *Dendrobium nobile* is caused by a deviation from the rule that one of the IRs is a reverse complement of the other; in that genome record, the two IRs represent a tandem repeat, which has artificially inflated the number of detected mismatches. In all other plastid genomes, the counts of IR mismatches was equal to, or less than, 55 nucleotides (Supplementary Table 2).

### Hypotheses on sequencing depth

The results of our statistical tests showed that the number of WRSD were significantly different across genome partitions as well as between the coding and non-coding sections of the plastid genomes. Specifically, a Kruskal-Wallis test identified that the distributions of the number of WRSD were significantly different across the four structural partitions of the plastid genome (*p* < 0.001; Table 3), and that the differences exhibited a large effect size (*d_c_* = 1.85). The pairwise comparison of the four partitions through post-hoc Wilcoxon rank-sum tests indicated that all states showed significant differences between their WRSD distributions (Table 4). These differences consistently exhibited a large effect size, except for the comparsion of IR_A_ against IR_B_ and of IR_A_ against the SSC, which exhibited small effect sizes (*d_c_* = 0.384 and *d_c_* = 0.435, respectively). Similarly, a Wilcoxon rank-sum test identified that the distribution of the number of WRSD was significantly different between the coding and non-coding sections of the plastid genomes (*p* < 0.001; Table 4) and that the difference exhibited a large effect size (*d_c_* = 2.169). Given these results, we reject hypotheses (a) and (b) and conclude that both the four structural partitions as well as the coding and non-coding sections of the plastid genomes exhibit significant differences in their number of WRSD and, by extension, their sequencing depth.

**Table 3.** P-values and effect sizes of the Kruskal-Wallis tests. Significance indicators: * *p* < 0.05, ** *p* < 0.01, *** *p* < 0.001. Effect size indicators: ^s^ small, ^m^ moderate, ^l^ large. Abbreviations are as in Table 2.

|  | Variable | n | p | d<sub>c</sub> |
| --- | --- | --- | --- | --- |
| Seq. depth | WRSD/partition | 664 | <0.001 *** | 1.850 <sup>l</sup> |
| Seq. evenness | Seq. platform | 187 | <0.001 *** | 0.707 <sup>m</sup> |
|  | Asm. software | 54 | 0.745 | 0.418 |

**Table 4.**
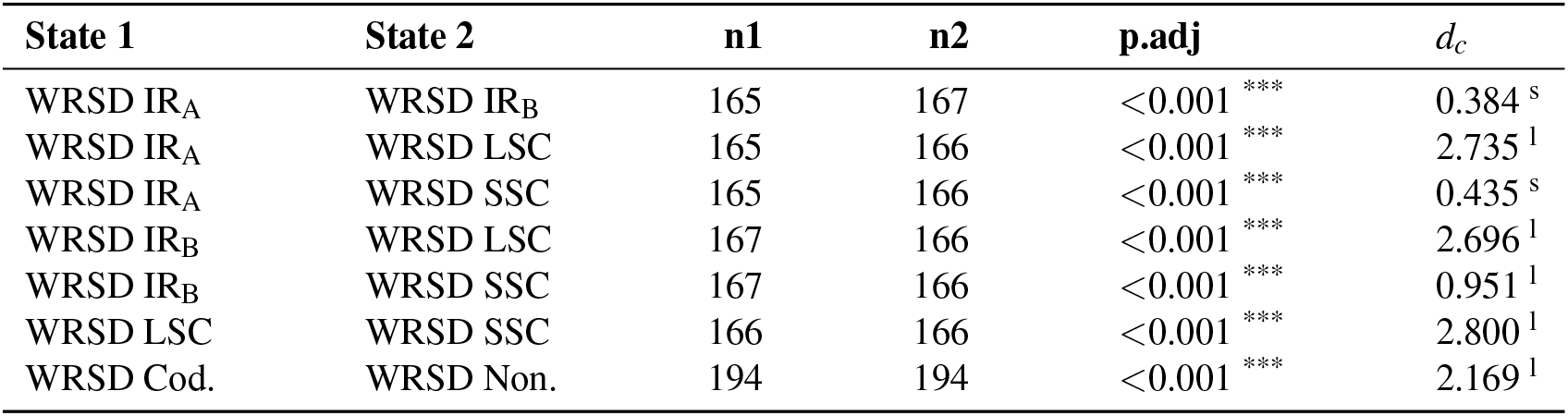
P-values and effect size of pairwise Wilcoxon rank-sum tests for the evaluation of differences in sequencing depth. All abbreviations are as in Table 1, all significance and effect size indicators as in Table 3. The p-value for the test between coding and non-coding genome sections did not undergo a Benjamini-Hochberg correction.

| State 1 | State 2 | n1 | n2 | p.adj | d<sub>c</sub> |
| --- | --- | --- | --- | --- | --- |
| WRSD IR<sub>A</sub> | WRSD IR<sub>B</sub> | 165 | 167 | <0.001 *** | 0.384 <sup>s</sup> |
| WRSD IR<sub>A</sub> | WRSD LSC | 165 | 166 | <0.001 *** | 2.735 <sup>l</sup> |
| WRSD IR<sub>A</sub> | WRSD SSC | 165 | 166 | <0.001 *** | 0.435 <sup>s</sup> |
| WRSD IR<sub>B</sub> | WRSD LSC | 167 | 166 | <0.001 *** | 2.696 <sup>l</sup> |
| WRSD IR<sub>B</sub> | WRSD SSC | 167 | 166 | <0.001 *** | 0.951 <sup>l</sup> |
| WRSD LSC | WRSD SSC | 166 | 166 | <0.001 *** | 2.800 <sup>l</sup> |
| WRSD Cod. | WRSD Non. | 194 | 194 | <0.001 *** | 2.169 <sup>l</sup> |

### Hypotheses on sequencing evenness

The results of our statistical tests showed that sequencing evenness in a plastid genome was significantly correlated with the number of its ambiguous nucleotides but not with the number of sequence mismatches between its two IRs. Specifically, Spearman rank correlation tests identified positive correlations between the genome-wide E-score and each of the two assembly quality metrics evaluated, but only the correlation between the E-score and the number of ambiguous nucleotides per genome was significant (*R_s_* = 0.14, *p* = 0.049; Figure 3), whereas the correlation between the E-score and the number of IR sequence mismatches was insignificant (*R_s_* = 0.12, *p* = 0.117). Given these results, we reject hypotheses (c) but not hypothesis (d) and conclude that the genome-wide E-score and, by extension, the sequencing evenness a plastid genome is correlated with the number of its ambiguous nucleotides.

**Figure 3.**
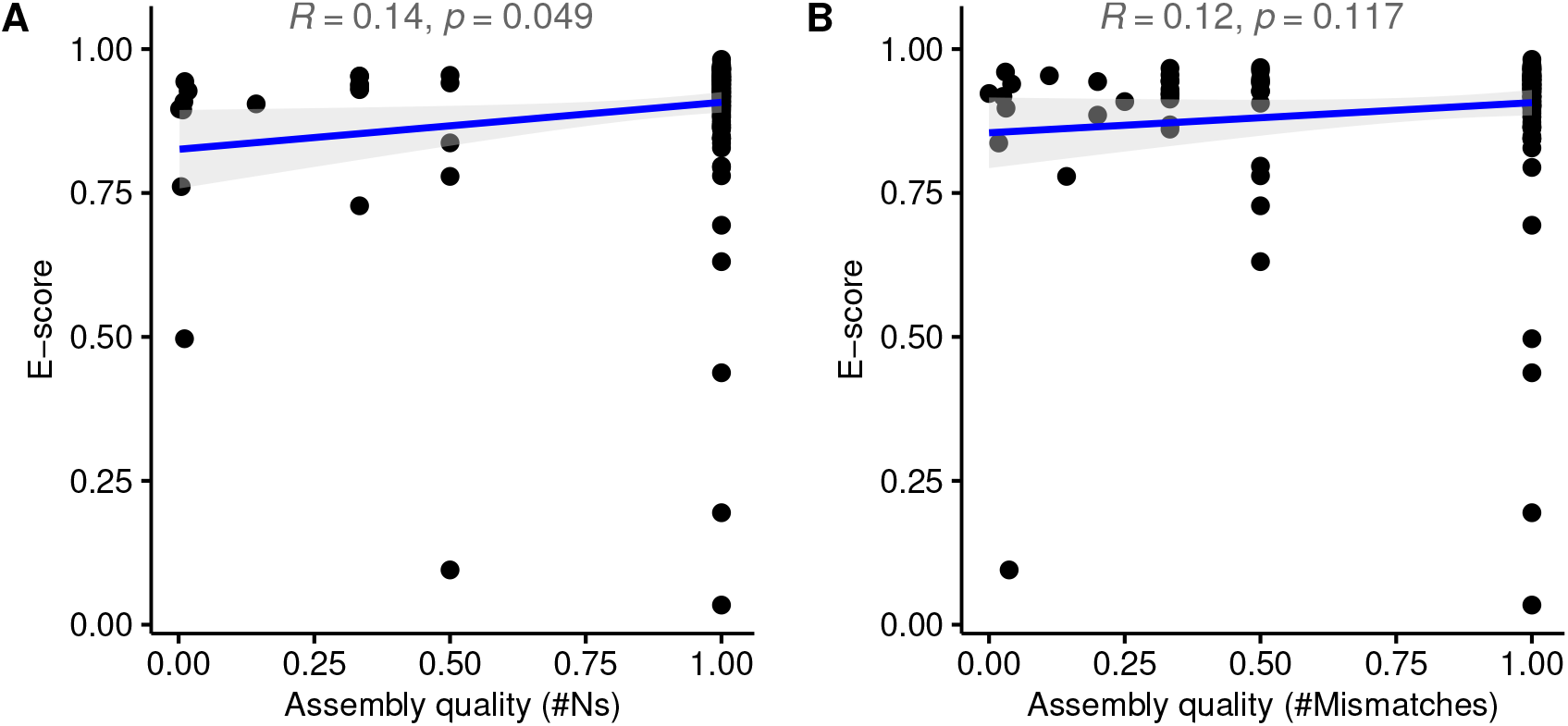
Regression plots to illustrate the correlation of the genome-wide E-score a plastid genome to (A) the number of ambiguous nucleotides per genome and (B) the number of sequence mismatches between its two IRs. The data values of each assembly quality metric were transformed to fit a range between 0 and 1. In each plot, the regression line is indicated in blue and its confidence interval through gray shades.

The results of our statistical tests also showed that sequencing evenness in plastid genomes was significantly different across sequencing platforms but not across assembly software. Specifically, a Kruskal-Wallis test identified that the genome-wide E-scores of plastid genomes were significantly different across sequencing platforms (*p* < 0.001; Table 3), and that these differences exhibited a moderate effect size (*d_c_* = 0.707). A graphical comparison of the data dispersion of the E-scores across different sequencing platforms indicated that HiSeq 1500 had a much narrower value dispersion than most of the other platforms (Figure 1C). Pairwise comparisons of the E-scores between sequencing platforms through post-hoc Wilcoxon rank-sum tests further indicated that ten (32%) of the 28 possible comparisons were statistically significant (Table 5). Large effect sizes were found for the differences between the Illumina platforms Genome Analyzer II and HiSeq 1500 (*d_c_* = 2.198), HiSeq 1500 and HiSeq 4000 (*d_c_* = 1.352), HiSeq 1500 and NextSeq 500 (*d_c_* = 3.000), and between HiSeq 2500 and NextSeq 500 (*d_c_* = 0.941). By contrast, the different assembly software employed to generate the plastid genomes did not result in significantly different E-scores of these genomes (*p* = 0.745; Table 3). A graphical comparison of the data dispersion of the E-scores across different assembly software indicated that only the software Velvet had a slightly narrower value dispersion than the other software tools evaluated (Figure 1D). Given these results, we reject hypotheses (e) but not hypothesis (f) and conclude that the genome-wide E-score and, by extension, the sequencing evenness a plastid genome is different across sequencing platforms.

**Table 5.**
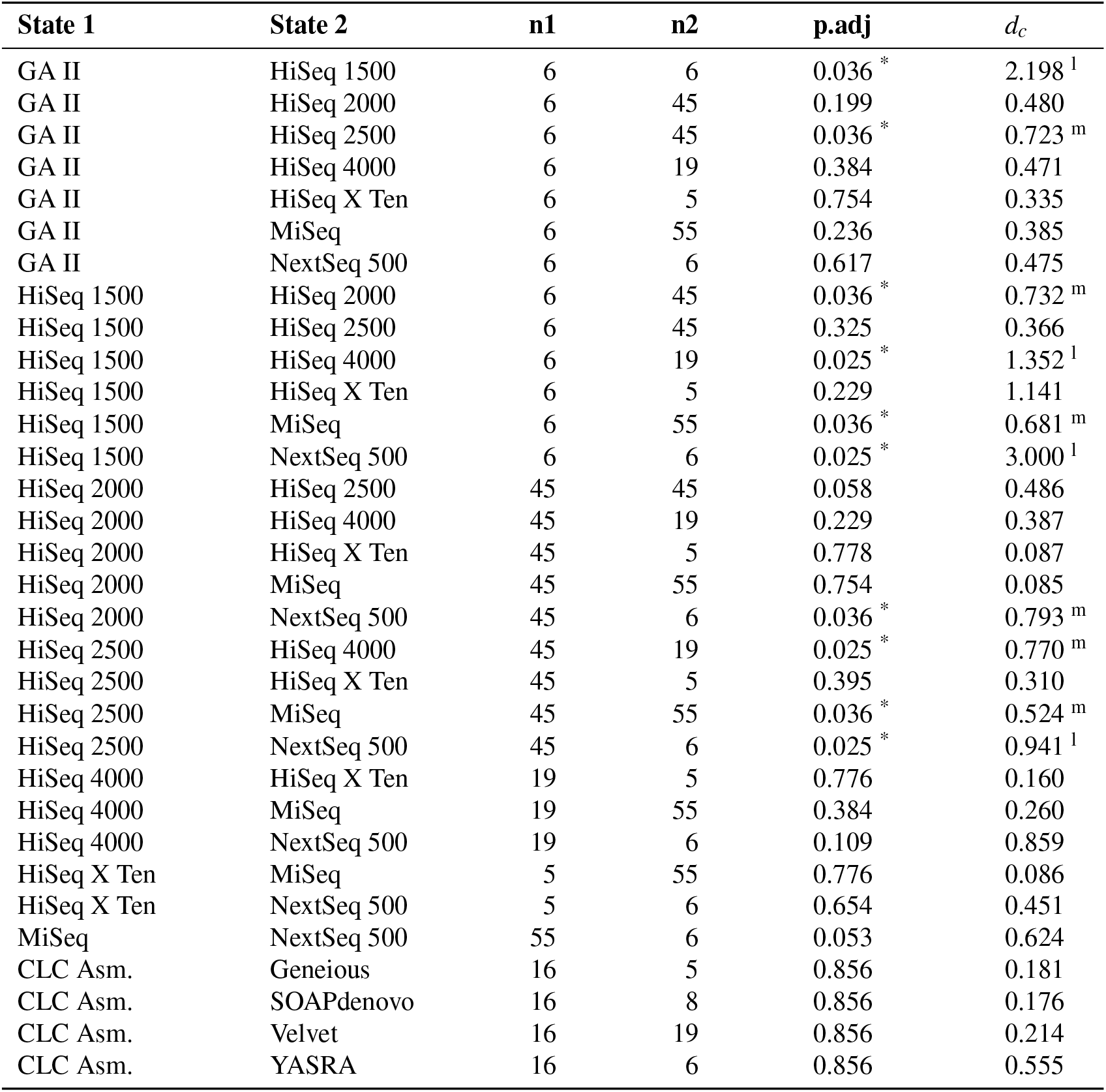
P-values and effect size of pairwise Wilcoxon rank-sum tests for the evaluation of differences in sequencing evenness. Prefix removal and abbreviations are as in Table 1, all significance and effect size indicators as in Table 3.

## DISCUSSION

### Sequencing evenness and assembly quality

Our results provide preliminary evidence for the conclusion that the observed correlation of sequencing evenness with assembly quality is not a product of chance alone. Specifically, we found a small but statistically significant correlation between the E-score and the number of ambiguous nucleotides per genome for the plastid genomes under study. The extent to which this finding can be generalized to a relevant correlation between sequencing evenness and assembly quality is dependent on the precision of the number of ambiguous nucleotides as an assembly quality indicator. In many studies, the quality of a genome assembly is evaluated through one or more indicators that can be measured across multiple genomes (Narzisi and Mishra, 2011; Gurevich et al., 2013). Several such indicators have been proposed and employed over time (e.g., Phillippy et al., 2008; Meader et al., 2010; Gurevich et al., 2013; Ou et al., 2018; Yang et al., 2019; Manchanda et al., 2020), and Smits (2019) even suggested the specification of a mandatory, standardized quality score for all genome records on NCBI Nucleotide to ensure an objective comparison of the stored sequences. In general, two groups of assembly quality indicators can be distinguished: alignment- and sequence-based indicators (Udall and Dawe, 2018). Indicators based on sequence alignments compare the synteny of nucleotides or functional annotations between genomes and can often be applied to partial genome sequences as well. Sequence-based indicators, by contrast, focus on sequence features such as length and contiguity and are typically agnostic to genome structure (Narzisi and Mishra, 2011; Udall and Dawe, 2018). In this investigation, we utilized both, a sequence-based indicator (i.e., the number of ambiguous nucleotides per genome) and an alignment-based indicator (i.e., the number of mismatches between the two plastid IRs), to evaluate assembly quality. The number of ambiguous nucleotides per genome has been employed as assembly quality indicator in various genomic investigations (e.g., Kuzmin et al., 2019; Yavas et al., 2019). The use of IR equity to indicate assembly quality is less common but has been suggested in several studies (Gruenstaeudl and Jenke, 2020; Mehl and Gruenstaeudl, 2020; Zheng et al., 2020). In fact, IR equity represents an easily accessible metric based on the quadripartite structure of plastid genomes (Jansen and Ruhlmann, 2012), reflects the natural repeat homogenization process in plastids (Goulding et al., 1996; Ruhlman et al., 2017), and is explicitly assumed by several software tools for the identification and visualization of plastid IRs (e.g., Lohse et al., 2013; Zheng et al., 2020). Our statistical analyses found that both indicators had a positive correlation with assembly quality in the plastid genomes under study, but that only the number of ambiguous nucleotides per genome was significantly correlated. This observation can be explained by the circumstance that many mismatches between the IRs of a plastid genome represent annotation errors rather than genuine sequence differences (Mehl and Gruenstaeudl, 2020) and, thus, do not typically impact sequencing depth. Ambiguous nucleotides of a genome sequence, by contrast, do impact the calculation of sequencing depth, rendering this indicator more sensitive to changes in sequencing evenness.

### Sequencing depth and genome structure

Our results provide preliminary evidence for the conclusion that differences in sequencing depth between the structural partitions of a plastid genome are not a product of chance alone. Specifically, we found that the number of WRSD was significantly different across the LSC, IR_B_, SSC, and IR_A_ of the plastid genomes under study. The extent to which this finding can be generalized to a relevant difference in sequencing depth between the four partitions of quadripartite plastid genomes is dependent on the importance of the length differences between the partitions. In most plastid genomes, the LSC is the longest of the four partitions and should – *ceteris paribus* – exhibit more WRSD on average than the other genome partitions. Indeed, the number of WRSD detected in the LSC was generally higher than those of the SSC or the IRs (Figure 1A), and the differences were significant and exhibited a large effect size in all comparisons involving the LSC (Table 4). However, significant differences in sequencing depth, albeit with small effect size, were also identified between the two IRs, which are identical in length. By default, the IRs of a plastid genome are homogenized in both length and sequence through recombination-dependent replication and natural gene conversion (Goulding et al., 1996; Ruhlman et al., 2017); non-identical IRs are only rarely observed (e.g., Turmel et al., 2017). The number of WRSD should, thus, be highly similar, if not identical, between the two IRs of a plastid genome. The significant differences in the number of WRSD between the four structural genome partitions may, thus, be indicative of length-independent differences in sequencing depth. Alternatively, the observed differences may represent a methodological artifact. For example, repeat regions are known to affect the bioinformatic process of mapping sequence reads to a reference genome (Schbath et al., 2012; Thankaswamy-Kosalai et al., 2017; Wang et al., 2021). In Bowtie2, for example, reads are mapped to a reference genome based on a mapping score that is identical across exact repeats, but pseudo-random numbers are used to break ties between repeat locations (Langmead and Salzberg, 2012). This stochasticity may result in unequal sequence coverages of the repeats if the overall sequencing depth is low (Straub et al., 2012). The border area of repeat regions remains a particular challenge for read mapping software, as reads span across the junction sites to non-repeat areas. The IRs of the plastid genome may, thus, exhibit different sequencing depth profiles despite equal sequence and length (Gruenstaeudl and Jenke, 2020). The results of this study support the conclusion of length-independent differences in sequencing depth, as numerous plastid genomes under study exhibited large differences in the number of WRSD between their IRs (Supplementary Table 2).

### Sequencing depth and coding status

Our results provide preliminary evidence for the conclusion that differences in sequencing depth between the coding and non-coding sections of a plastid genome are not a product of chance alone. Specifically, we found a significant difference in the number of WRSD between coding and non-coding sections of the plastid genomes under study as well as a large effect size for this difference. Both the mean and the median of the number of WRSD in the coding sections were larger than their counterparts in the non-coding sections (Table 2). This difference is, thus, unlikely the result of the different lengths of the two sections, as the cumulative length of all coding regions in a plastid genome is typically smaller than the cumulative length of all non-coding regions. Instead, the difference in the number of WRSD between coding and non-coding genome sections may be the result of different levels of GC content (Romiguier and Roux, 2017). GC content is known to influence the rate of sequencing error in genome sequences (often termed ‘GC bias’; Benjamini and Speed, 2012; Dabney and Meyer, 2012; Chen et al., 2013) and can also affect the sequence coverage of a genome assembly (Rieber et al., 2013). Plastid genomes are known to exhibit uneven GC distributions across their sequences (e.g., Li et al., 2016; Mower et al., 2019). The observed differences in sequencing depth between the coding and non-coding genome sections may, thus, be related to the varying levels of GC content.

### Sequencing evenness and sequencing platform

Our results provide preliminary evidence for the conclusion that differences in sequencing evenness between different sequencing platforms are not a product of chance alone. Specifically, we found a significant difference of the E-score values of different Illumina sequencing platforms as well as a moderate effect size for this difference among the plastid genomes under study (Table 3). Post-hoc test revealed significant differences in, and moderate to large effect sizes for, several pairwise comparisons, particularly those involving the sequencing platforms HiSeq1500 and NextSeq 500 (Table 5). These results may be indicative of the different error rates of the various NGS sequencing platforms (Fox et al., 2014; Pfeiffer et al., 2018), which may become manifest as different sequencing depth profiles. In fact, several studies have reported differences in the sequencing depth of plastid genomes that were sequenced on different sequencing platforms (e.g., Brozynska et al., 2014; Wu et al., 2014). Additional studies are warranted to evaluate the precise impact that the selection of sequencing platform has on the depth and evenness of sequence coverage of plastid genomes.

### Sequencing evenness and software choice

No evidence for a significant difference in sequencing evenness between different assembly software tools was detected among the plastid genomes under study. This result is in opposition to several previous studies that report a considerable impact by the choice of assembly software on the quality of the assembled plastid genomes. Dierckxsens et al. (2016), for example, found considerable differences in the number of plastid contigs, the sequence coverage of these contigs, and in the assembly accuracy of five different assembly software tools. Similarly Freudenthal et al. (2020) compared different software tools for plastid genome assembly and found that the tools differed in their accuracy, among other factors; sequence coverage was hereby employed as one of several components in a special metric to evaluate the accuracy of plastid genome assemblies. Evidently, the choice of assembly software can have a considerable impact on the assembly results, even if sequencing evenness may not be affected at a significant level.

## DECLARATIONS

### Author Contributions

NJ - Implementation, documentation, testing, and manuscript writing. MG - Conception, project design, oversight, implementation, documentation, and manuscript writing. Both authors have read and approved the final version of the manuscript.

## Acknowledgments

The authors acknowledge the high-performance computing service of the ZEDAT of the Freie Universität Berlin for providing allocations of computing time. A part of this research was conducted by N.J. toward a master of science degree.

## Data Availability Statement

The bioinformatic pipeline designed and employed to conduct the data retrieval, parsing, and analysis steps of this investigation is archived on GitHub at https://github.com/nilsj9/PlastidSequenceCoverage. All data sets generated and analyzed are archived on Zenodo at https://doi.org/10.5281/zenodo.4555956.

## Supplemental Material

Supplementary Tables 1 and 2 as well as Supplementary Figures 1–13 are available as online supplements.

