## Supplemental Material for "Depth and evenness of sequence coverage are associated with assembly quality, genome structure, and choice of sequencing platform in archived plastid genomes"

**Supplementary Table 1:** Overview of the species names, taxonomic associations, and genome record accession numbers (NCBI Nucleotide and NCBI SRA) of all plastid genome under study.

| Order | Family | Species | NCBI Nucl. | NCBI SRA |
|---|---|---|---|---|
| | | **ANGIOSPERMS** | | |
| | | **Angiosperms – Eudicots – Rosids** | | |
| Asterales | Asteraceae | *Artemisia annua* | NC_034683 | SRR5602595 |
| | | *Echinacea angustifolia* | NC_034324 | SRR5602579 |
| | | *Echinacea atrorubens* | NC_034323 | SRR5602578 |
| | | *Echinacea laevigata* | NC_034322 | SRR5602581 |
| | | *Echinacea pallida* | NC_034321 | SRR5602580 |
| | | *Echinacea paradoxa* | NC_034320 | SRR5602575 |
| | | *Echinacea purpurea* | NC_034327 | SRR5602574 |
| | | *Echinacea sanguinea* | NC_034328 | SRR5602577 |
| | | *Echinacea speciosa* | NC_034325 | SRR5602576 |
| | | *Echinacea tennesseensis* | NC_034326 | SRR5602587 |
| | | *Helianthus annuus* | NC_007977 | SRR825830 |
| | | *Helianthus argophyllus* | NC_030275 | SRR2155086 |
| | | *Helianthus debilis* | NC_030173 | SRR5907791 |
| | | *Carthamus tinctorius* | NC_030783 | SRR2154065 |
| | | *Centaurea diffusa* | NC_024286 | SRR2729212 |
| | | *Taraxacum kok-saghyz* | NC_032057 | SRR8185394 |
| | | *Lactuca sativa* | NC_007578 | SRR8736654 |
| Ericales | Actinidiaceae | *Actinidia chinensis* | NC_026690 | DRR083750 |
| | Ebenaceae | *Diospyros lotus* | NC_030786 | SRR1560932 |
| | Ericaceae | *Vaccinium macrocarpon* | NC_019616 | SRR1276173 |
| | Primulaceae | *Primula veris* | NC_031428 | SRR1660449 |
| Gentianales | Rubiaceae | *Mitragyna speciosa* | NC_034698 | SRR5602600 |
| | | *Coffea canephora* | NC_030053 | ERR321701 |
| Lamiales | Lamiaceae | *Premna microphylla* | NC_026291 | SRR6940036 |
| | | *Tectona grandis* | NC_020098 | SRR6940065 |
| | | *Haplostachys haplostachya* | NC_029819 | SRR3170741 |
| | | *Stachys byzantina* | NC_029825 | SRR3170744 |
| | | *Stachys chamissonis* | NC_029822 | SRR3170745 |
| | | *Stachys coccinea* | NC_029823 | SRR3170746 |
| | | *Stachys sylvatica* | NC_029824 | SRR3170747 |
| | | *Perilla frutescens* | NC_030756 | SRR6940083 |
| | | *Ocimum basilicum* | NC_035143 | SRR6940087 |
| | | *Scutellaria baicalensis* | NC_027262 | SRR6940088 |
| | | *Ajuga reptans* | NC_023102 | SRR6940062 |
| | Oleaceae | *Jasminum sambac* | NC_034694 | SRR5602611 |
| | | *Jasminum tortuosum* | NC_034691 | SRR5602601 |
| | | *Olea europaea* subsp. *europaea* | NC_015401 | ERR375848 |
| | Phrymaceae | *Erythranthe lutea* | NC_030212 | SRR5307620 |
| | Plantaginaceae | *Digitalis lanata* | NC_034688 | SRR5602573 |
| | Verbenaceae | *Aloysia citrodora* | NC_034695 | SRR5602597 |
| Solanales | Convolvulaceae | *Ipomoea batatas* | NC_026703 | SRR7868503 |
| | | *Ipomoea trifida* | NC_034670 | SRR6667669 |
| | Solanaceae | *Nicotiana otophora* | NC_032724 | SRR1171700 |
| | | *Solanum berthaultii* | NC_034951 | SRR5349620 |
| | | *Solanum commersonii* | NC_028069 | SRR5349609 |
| | | *Solanum melongena* | NC_030207 | DRR014074 |
| | | **Angiosperms – Eudicots – Asterids** | | |
| Brassicales | Brassicaceae | *Arabis alpina* | NC_023367 | SRR7880736 |

Continued on next page

Supplementary Table 1 – continued from previous page

| Order | Family | Species | NCBI Nucl. | NCBI SRA |
|---|---|---|---|---|
| | | *Brassica napus* | NC_016734 | SRR6856349 |
| | | *Brassica nigra* | NC_030450 | SRR2054762 |
| | | *Arabidopsis halleri* | NC_034366 | SRR8040824 |
| | | *Arabidopsis lyrata* | NC_034365 | SRR8157563 |
| | | *Arabidopsis thaliana* | NC_000932 | SRR3340908 |
| | | *Capsella bursa-pastoris* | NC_009270 | SRR5412136 |
| | | *Capsella grandiflora* | NC_028517 | SRR1508428 |
| | | *Capsella rubella* | NC_027693 | ERR636124 |
| | | *Thlaspi arvense* | NC_034362 | SRR1034659 |
| Cucurbitales | Cucurbitaceae | *Cucumis melo* subsp. *melo* | NC_015983 | ERR246536 |
| Fabales | Fabaceae | *Cicer arietinum* | NC_011163 | SRR5183095 |
| | | *Vicia sativa* | NC_027155 | ERR413103 |
| | | *Glycyrrhiza glabra* | NC_024038 | SRR8690419 |
| | | *Lotus japonicus* | NC_002694 | SRR8115522 |
| | | *Cajanus cajan* | NC_031429 | SRR073338 |
| | | *Glycine soja* | NC_022868 | SRR7725010 |
| | | *Phaseolus vulgaris* | NC_009259 | SRR5807695 |
| | | *Vigna unguiculata* | NC_018051 | SRR7125688 |
| | | *Wisteria floribunda* | NC_027677 | SRR1265941 |
| | Betulaceae | *Betula nana* | NC_033978 | ERR2026268 |
| | | *Ostrya rehderiana* | NC_028349 | SRR8302715 |
| | Juglandaceae | *Juglans regia* | NC_028617 | SRR2057822 |
| Geraniales | Francoaceae | *Melianthus villosus* | NC_023256 | SRR576532 |
| | Geraniaceae | *Pelargonium citronellum* | NC_031194 | SRR576534 |
| Malpighiales | Chrysobalanaceae | *Chrysobalanus icaco* | NC_024061 | SRR1179655 |
| | | *Couepia guianensis* | NC_024063 | SRR1179648 |
| | | *Hirtella physophora* | NC_024066 | SRR1179646 |
| | | *Hirtella racemosa* | NC_024060 | SRR1179649 |
| | | *Licania alba* | NC_024064 | SRR1179652 |
| | | *Licania sprucei* | NC_024065 | SRR1179651 |
| | | *Parinari campestris* | NC_024067 | SRR1179645 |
| | Euphorbiaceae | *Euphorbia esula* | NC_033910 | SRR6355713 |
| | Salicaceae | *Populus alba* | NC_008235 | SRR3678826 |
| | | *Populus tremula* | NC_027425 | ERR1735633 |
| | | *Populus tremula x alba* | NC_028504 | SRR1653109 |
| | | *Populus trichocarpa* | NC_009143 | SRR5467392 |
| Malvales | Cytinaceae | *Cytinus hypocistis* | NC_031150 | ERR964904 |
| Malvales | Malvaceae | *Abelmoschus esculentus* | NC_035234 | SRR5812498 |
| | | *Althaea officinalis* | NC_034701 | SRR5602596 |
| | | *Gossypium herbaceum* | NC_023215 | SRR2012721 |
| | | *Hibiscus syriacus* | NC_026909 | SRR1265942 |
| Myrtales | Lythraceae | *Punica granatum* | NC_035240 | SRR5812494 |
| | Myrtaceae | *Pimenta dioica* | NC_034684 | SRR5602585 |
| Rosales | Cannabaceae | *Cannabis sativa* | NC_026562 | SRR7285294 |
| | | *Humulus lupulus* | NC_028032 | SRR8690418 |
| | Moraceae | *Ficus carica* | NC_035237 | SRR5678803 |
| | | *Ficus racemosa* | NC_028185 | SRR1405699 |
| | Rosaceae | *Prunus dulcis* | NC_034696 | SRR5602582 |
| | | *Prunus kansuensis* | NC_023956 | SRR3138168 |
| | | *Prunus mume* | NC_023798 | SRR8240066 |
| | | *Prunus yedoensis* | NC_026980 | DRR169775 |
| | | *Malus prunifolia* | NC_031163 | SRR3571175 |
| | | *Fragaria chiloensis* | NC_019601 | SRR1612828 |
| | | *Fragaria vesca subsp. bracteata* | NC_018766 | SRR866156 |
| | | *Fragaria virginiana* | NC_019602 | SRR5602605 |
| Sapindales | Anacardiaceae | *Anacardium occidentale* | NC_035235 | SRR5812497 |

Continued on next page

Supplementary Table 1 – continued from previous page

| Order | Family | Species | NCBI Nucl. | NCBI SRA |
|---|---|---|---|---|
| | | *Mangifera indica* | NC_035239 | SRR5812496 |
| | | *Pistacia vera* | NC_034998 | SRR4453367 |
| | Burseraceae | *Boswellia sacra* | NC_029420 | SRR5602594 |
| | Rutaceae | *Citrus limon* | NC_034690 | SRR5602593 |
| | Sapindaceae | *Litchi chinensis* | NC_035238 | SRR5812499 |
| Vitales | Vitaceae | *Vitis amurensis* | NC_031383 | SRR5891950 |
| | | *Vitis rotundifolia* | NC_023790 | SRR5627788 |
| | | **Angiosperms – Eudicots – Other** | | |
| Caryophyllales | Cactaceae | *Carnegiea gigantea* | NC_027618 | SRR5036293 |
| | Chenopodiaceae | *Chenopodium quinoa* | NC_034949 | SRR5938314 |
| | Droseraceae | *Aldrovanda vesiculosa* | NC_035416 | SRR7072768 |
| | | *Dionaea muscipula* | NC_035417 | SRR7072324 |
| | | *Drosera erythrorhiza* | NC_035241 | SRR7072766 |
| | | *Drosera regia* | NC_035415 | SRR7072322 |
| | Polygonaceae | *Fagopyrum tataricum* | NC_027161 | SRR5433722 |
| Ranunculales | Berberidaceae | *Epimedium koreanum* | NC_029943 | SRR8534306 |
| | Ranunculaceae | *Hydrastis canadensis* | NC_034702 | SRR5602606 |
| | | **Angiosperms – Monodicots** | | |
| Arecales | Arecaceae | *Elaeis guineensis* | NC_017602 | ERR276848 |
| | | *Podococcus barteri* | NC_027276 | SRR2120220 |
| | | *Phoenix dactylifera* | NC_013991 | SRR2518264 |
| Asparagales | Amaryllidaceae | *Allium cepa* | NC_024813 | SRR1686960 |
| | Asparagaceae | *Asparagus officinalis* | NC_034777 | DRR056675 |
| | Orchidaceae | *Apostasia odorata* | NC_030722 | SRR6037860 |
| | | *Cymbidium ensifolium* | NC_028525 | SRR6117755 |
| | | *Cymbidium kanran* | NC_029711 | SRR6117756 |
| | | *Cymbidium lancifolium* | NC_029712 | SRR6117757 |
| | | *Cymbidium macrorhizon* | NC_029713 | SRR6117754 |
| | | *Dendrobium aphyllum* | NC_035322 | SRR3932123 |
| | | *Dendrobium chrysanthum* | NC_035336 | SRR3932147 |
| | | *Dendrobium chrysotoxum* | NC_028549 | SRR3932141 |
| | | *Dendrobium crepidatum* | NC_035331 | SRR3932124 |
| | | *Dendrobium denneanum* | NC_035324 | SRR3932144 |
| | | *Dendrobium falconeri* | NC_035326 | SRR3932139 |
| | | *Dendrobium nobile* | NC_029456 | SRR3932122 |
| | | *Dendrobium parishii* | NC_035339 | SRR3932128 |
| | | *Dendrobium pendulum* | NC_029705 | SRR3932138 |
| | | *Dendrobium primulinum* | NC_035321 | SRR3932120 |
| | | *Dendrobium wardianum* | NC_035329 | SRR3932135 |
| | | *Ludisia discolor* | NC_030540 | SRR3484539 |
| Dioscoreales | Dioscoreaceae | *Dioscorea rotundata* | NC_024170 | DRR063110 |
| | | *Dioscorea villosa* | NC_034686 | SRR5602590 |
| Liliales | Liliaceae | *Lilium tsingtauense* | NC_027675 | SRR1265940 |
| Poales | Poaceae | *Phyllostachys edulis* | NC_015817 | SRR8245858 |
| | | *Oryza barthii* | NC_027460 | SRR7341625 |
| | | *Oryza glumipatula* | NC_027461 | DRR057991 |
| | | *Oryza longiglumis* | NC_034763 | DRR056658 |
| | | *Oryza longistaminata* | NC_027462 | DRR058031 |
| | | *Oryza meridionalis* | NC_016927 | DRR058012 |
| | | *Oryza meyeriana* | NC_034765 | DRR056670 |
| | | *Oryza officinalis* | NC_027463 | DRR000604 |
| | | *Oryza punctata* | NC_027676 | SRR1264539 |
| | | *Oryza ridleyi* | NC_034764 | DRR056663 |
| | | *Oryza rufipogon* | NC_017835 | SRR6220521 |

Continued on next page

Supplementary Table 1 – continued from previous page

| Order | Family | Species | NCBI Nucl. | NCBI SRA |
|---|---|---|---|---|
| | | *Oryza sativa* | NC_031333 | ERR2696318 |
| | | *Avena sativa* | NC_027468 | SRR6056489 |
| | | *Dactylis glomerata* | NC_027473 | SRR5236602 |
| | | *Deschampsia antarctica* | NC_023533 | SRR1158316 |
| | | *Aegilops sharonensis* | NC_024816 | ERR359708 |
| | | *Arundo plinii* | NC_034652 | SRR4319202 |
| | | *Eragrostis tef* | NC_029413 | SRR1463402 |
| | | *Sporobolus michauxianus* | NC_029416 | SRR4434178 |
| | | *Miscanthus sacchariflorus* | NC_028720 | SRR559245 |
| | | *Miscanthus sinensis* | NC_028721 | SRR4028761 |
| | | *Sorghum timorense* | NC_023800 | SRR424217 |
| | | *Alloteropsis angusta* | NC_027951 | SRR7528995 |
| | | *Alloteropsis cimicina* | NC_027952 | SRR7529015 |
| | | *Alloteropsis paniculata* | NC_032078 | SRR4051980 |
| | | *Alloteropsis semialata* | NC_027824 | SRR7528994 |
| | | *Echinochloa crus-galli* | NC_028719 | SRR5920285 |
| | | *Echinochloa oryzicola* | NC_024643 | SRR5902661 |
| | | *Cenchrus americanus* | NC_024171 | SRR5204424 |
| | | *Panicum capillare* | NC_030493 | SRR485871 |
| | | **Angiosperms – Basal Angiosperms** | | |
| Austrobaileyales | Schisandraceae | *Illicium anisatum* | NC_034703 | SRR5602608 |
| | | *Illicium floridanum* | NC_034685 | SRR5602609 |
| | | *Illicium henryi* | NC_034699 | SRR5602610 |
| | | *Illicium verum* | NC_034689 | SRR5602607 |
| Laurales | Lauraceae | *Cinnamomum verum* | NC_035236 | SRR5812493 |
| | | *Laurus nobilis* | NC_034700 | SRR5602602 |
| Magnoliales | Magnoliaceae | *Magnolia biondii* | NC_034687 | SRR5602588 |
| Piperales | Piperaceae | *Piper auritum* | NC_034697 | SRR5602592 |
| | | *Piper nigrum* | NC_034692 | SRR5602591 |
| | | **GYMNOSPERMS** | | |
| Cupressales | Cupressaceae | *Juniperus cedrus* | NC_028190 | SRR1145775 |
| | | *Juniperus communis* | NC_035068 | ERR268423 |
| | Taxaceae | *Taxus baccata* | NC_035066 | ERR268425 |
| Gnetales | Gnetaceae | *Gnetum gnemon* | NC_026301 | ERR268420 |
| Pinales | Pinaceae | *Picea glauca* | NC_028594 | SRR869482 |
| | | *Picea abies* | NC_021456 | ERR1727007 |
| | | *Picea asperata* | NC_032367 | ERR1735493 |
| | | *Picea crassifolia* | NC_032366 | ERR1735613 |
| | | *Pinus massoniana* | NC_021439 | SRR7666034 |
| | | *Pinus sylvestris* | NC_035069 | ERR268429 |
| | | *Pinus taeda* | NC_021440 | SRR1049756 |

**Supplementary Table 2:** Overview of the metadata, the E-score value, the number of WRSD per genome partition and coding status, and the assembly quality metrics of each plastid genome under study. Dashes indicate values that were either missing from the genome records or could not be parsed through the methods described. Abbreviations used: Asm. = Assembler/Assembly; Cod. = Coding; GA = Genome Analyzer; Mism. = Mismatch; Non. = Non-coding; Seq. = Sequencing.

| Sample | Metadata | | E-score | WRSD | | | | | | Asm. quality | |
|---|---|---|---|---|---|---|---|---|---|---|---|
| | Seq. platform | Asm. software | | LSC | $IR_B$ | SSC | $IR_A$ | Cod. | Non. | Ns | Mism. |
| NC_035234 | HiSeq 2500 | Velvet | 0.92 | - | - | - | - | 50 | 31 | 0 | - |
| NC_026690 | HiSeq 4000 | Velvet | 0.95 | 55 | 18 | 15 | 13 | 54 | 14 | 0 | 0 |
| NC_024816 | HiSeq 2000 | Consed | 0.95 | 52 | 14 | 7 | 5 | 26 | 30 | 0 | 0 |
| NC_023102 | HiSeq 2500 | Velvet | 0.95 | 41 | 12 | 10 | 8 | 50 | 3 | 0 | 0 |
| NC_035416 | HiSeq 1500 | - | 0.95 | 48 | 21 | 7 | 18 | 57 | 16 | 0 | 0 |
| NC_024813 | HiSeq 2000 | Newbler Asm. | 0.92 | 54 | 24 | 13 | 23 | 54 | 13 | 0 | 0 |
| NC_027951 | HiSeq 2500 | - | 0.96 | 43 | 9 | 2 | 2 | 34 | 13 | 0 | 0 |
| NC_027952 | HiSeq 2500 | - | 0.97 | 32 | 6 | 9 | 4 | 19 | 13 | 0 | 1 |
| NC_032078 | HiSeq 2500 | - | 0.95 | 55 | 1 | 8 | 7 | 37 | 18 | 0 | 8 |
| NC_027824 | HiSeq 2500 | - | 0.96 | 18 | 8 | 4 | 5 | 23 | 25 | 0 | 1 |
| NC_034695 | MiSeq | - | 0.94 | 50 | 23 | 11 | 19 | 73 | 9 | 2 | 0 |
| NC_034701 | MiSeq | - | 0.94 | 51 | 15 | 15 | 14 | 61 | 12 | 0 | 0 |
| NC_035235 | HiSeq 2500 | Velvet | 0.95 | 50 | 23 | 14 | 24 | 55 | 32 | 0 | 0 |
| NC_030722 | HiSeq 4000 | - | 0.86 | 55 | 14 | 13 | 15 | 55 | 29 | 0 | 0 |
| NC_034366 | HiSeq 2500 | - | 0.97 | 38 | 22 | 6 | 12 | 51 | 7 | 0 | 0 |
| NC_034365 | HiSeq 4000 | - | 0.91 | 55 | 20 | 13 | 17 | 59 | 19 | 0 | 1 |
| NC_000932 | HiSeq 1500 | - | 0.97 | - | - | - | - | 49 | 10 | 0 | - |
| NC_023367 | HiSeq 2500 | - | 0.89 | 57 | 20 | 11 | 15 | 40 | 36 | 0 | 0 |
| NC_034683 | MiSeq | - | 0.92 | 57 | 14 | 10 | 10 | 50 | 12 | 0 | 0 |
| NC_034652 | MiSeq | SPAdes | 0.80 | 52 | 11 | 4 | 10 | 7 | 20 | 0 | 1 |
| NC_034777 | HiSeq 2500 | - | 0.98 | 26 | 16 | 5 | 1 | 58 | 13 | 0 | 0 |
| NC_027468 | MiSeq | Velvet | 0.97 | 40 | 8 | 7 | 1 | 25 | 11 | 0 | 0 |
| NC_033978 | NextSeq 500 | - | 0.90 | - | 25 | - | - | 43 | 31 | 0 | - |
| NC_029420 | MiSeq | - | 0.92 | 35 | 22 | 11 | 20 | 55 | 33 | 0 | 0 |
| NC_016734 | HiSeq 3000 | - | 0.78 | 51 | 21 | 12 | 20 | 38 | 19 | 0 | 1 |
| NC_030450 | HiSeq 2500 | CLC Asm. | 0.90 | 46 | 16 | 12 | 13 | 66 | 21 | 0 | 0 |
| NC_031429 | GA II | SPAdes | 0.84 | 47 | 11 | 5 | 11 | 60 | 24 | 0 | 0 |
| NC_026562 | NextSeq 500 | SOAPdenovo | 0.63 | 19 | 8 | 14 | 7 | 1 | 0 | 0 | 1 |
| NC_009270 | MiSeq | - | 0.94 | 45 | 24 | 11 | 18 | 55 | 24 | 0 | 0 |
| NC_028517 | HiSeq 2000 | CLC Asm. | 0.94 | 55 | 23 | 11 | 24 | 62 | 15 | 0 | 0 |
| NC_027693 | GA II | CLC Asm. | 0.91 | 45 | 24 | 13 | 23 | 52 | 21 | 0 | 0 |
| NC_027618 | MiSeq | SOAPdenovo | 0.91 | - | - | - | - | 53 | 20 | 0 | - |
| NC_030783 | HiSeq 1500 | CLC Asm. | 0.93 | 48 | 26 | 11 | 22 | 54 | 11 | 0 | 1 |
| NC_024171 | HiSeq 2500 | MIRA | 0.90 | 44 | 16 | 6 | 14 | 47 | 21 | 6 | 0 |
| NC_024286 | HiSeq 2000 | Ray | 0.97 | 44 | 5 | 18 | 9 | 55 | 24 | 0 | 2 |
| NC_034949 | HiSeq 2500 | CLC Asm. | 0.94 | 50 | 19 | 12 | 18 | 61 | 8 | 0 | 0 |
| NC_024061 | HiSeq 2000 | - | 0.94 | 51 | 24 | 11 | 20 | 63 | 11 | 0 | 0 |
| NC_011163 | HiSeq 2000 | - | 0.88 | - | - | - | - | 37 | 27 | 0 | - |
| NC_035236 | HiSeq 2500 | Velvet | 0.94 | 60 | 7 | 12 | 6 | 46 | 28 | 0 | 0 |
| NC_034690 | MiSeq | - | 0.95 | 50 | 16 | 12 | 12 | 49 | 14 | 2 | 0 |
| NC_030053 | - | Newbler Asm. | 0.92 | 55 | 16 | 11 | 15 | 44 | 11 | 0 | 38 |
| NC_024063 | HiSeq 2000 | - | 0.94 | 55 | 17 | 13 | 18 | 50 | 9 | 0 | 0 |
| NC_015983 | GA IIx | Newbler Asm. | 0.91 | 51 | 27 | 5 | 26 | 49 | 15 | 0 | 0 |
| NC_028525 | MiSeq | - | 0.90 | 45 | 6 | 18 | 6 | 35 | 24 | 0 | 0 |
| NC_029711 | MiSeq | Geneious | 0.96 | 51 | 17 | 10 | 17 | 59 | 11 | 0 | 0 |
| NC_029712 | MiSeq | Geneious | 0.96 | 42 | 13 | 9 | 11 | 67 | 9 | 0 | 0 |
| NC_029713 | MiSeq | Geneious | 0.44 | 0 | 0 | 4 | 0 | 0 | 0 | 0 | 0 |
| NC_031150 | HiSeq 2000 | OBITools | 0.88 | - | - | - | - | 8 | 2 | 0 | - |
| NC_027473 | HiSeq X Ten | Velvet | 0.89 | 54 | 14 | 9 | 11 | 42 | 21 | 0 | 0 |

Continued on next page

Supplementary Table 2 – continued from previous page

| Sample | Metadata | | E-score | WRSD | | | | | | Asm. quality | |
|---|---|---|---|---|---|---|---|---|---|---|---|
| | Seq. platform | Asm. software | | LSC | $IR_B$ | SSC | $IR_A$ | Cod. | Non. | Ns | Mism. |
| NC_035322 | HiSeq 4000 | - | 0.91 | 43 | 7 | 7 | 7 | 25 | 22 | 0 | 0 |
| NC_035336 | HiSeq 4000 | - | 0.92 | 38 | 6 | 5 | 7 | 30 | 28 | 0 | 0 |
| NC_028549 | HiSeq 4000 | - | 0.87 | 60 | 11 | 9 | 9 | 34 | 54 | 0 | 0 |
| NC_035331 | HiSeq 4000 | - | 0.91 | 46 | 10 | 4 | 7 | 32 | 30 | 0 | 2 |
| NC_035324 | HiSeq 4000 | - | 0.93 | 40 | 9 | 4 | 8 | 31 | 29 | 0 | 0 |
| NC_035326 | HiSeq 4000 | - | 0.91 | 49 | 4 | 5 | 7 | 26 | 29 | 0 | 0 |
| NC_029456 | HiSeq 4000 | - | 0.92 | 46 | 7 | 7 | 8 | 40 | 24 | 0 | 12830 |
| NC_035339 | HiSeq 4000 | - | 0.91 | 39 | 10 | 13 | 10 | 26 | 19 | 0 | 0 |
| NC_029705 | HiSeq 4000 | - | 0.89 | 47 | 9 | 12 | - | 43 | 33 | 0 | - |
| NC_035321 | HiSeq 4000 | - | 0.92 | 42 | 7 | 11 | 6 | 23 | 24 | 0 | 0 |
| NC_035329 | HiSeq 4000 | - | 0.87 | 58 | 10 | 10 | 10 | 41 | 33 | 0 | 2 |
| NC_023533 | HiSeq 2000 | - | 0.93 | 59 | 19 | 7 | 19 | 45 | 5 | 0 | 0 |
| NC_034688 | MiSeq | - | 0.94 | 46 | 16 | 11 | 14 | 63 | 9 | 0 | 0 |
| NC_035417 | HiSeq 1500 | - | 0.97 | - | - | - | - | 34 | 13 | 0 | - |
| NC_024170 | MiSeq | MIRA | 0.10 | 0 | 0 | 0 | 0 | 0 | 0 | 1 | 26 |
| NC_034686 | MiSeq | - | 0.94 | 53 | 18 | 12 | 18 | 65 | 15 | 0 | 0 |
| NC_030786 | HiSeq 2000 | FLASH | 0.93 | 44 | 9 | 14 | 13 | 46 | 27 | 0 | 1 |
| NC_035241 | HiSeq 1500 | - | 0.95 | 32 | 27 | 4 | 27 | 43 | 27 | 0 | 0 |
| NC_035415 | HiSeq 1500 | - | 0.97 | 31 | 14 | 6 | 10 | 47 | 11 | 0 | 0 |
| NC_034324 | MiSeq | - | 0.84 | 47 | 19 | 12 | 13 | 44 | 11 | 0 | 0 |
| NC_034323 | MiSeq | - | 0.92 | 47 | 18 | 13 | 13 | 66 | 9 | 0 | 0 |
| NC_034322 | MiSeq | - | 0.92 | 60 | 13 | 10 | 12 | 41 | 16 | 0 | 0 |
| NC_034321 | MiSeq | - | 0.79 | 47 | 21 | 15 | 12 | 64 | 17 | 0 | 0 |
| NC_034320 | MiSeq | - | 0.90 | 53 | 19 | 13 | 15 | 56 | 15 | 0 | 0 |
| NC_034327 | MiSeq | - | 0.88 | 57 | 18 | 11 | 15 | 46 | 20 | 0 | 0 |
| NC_034328 | MiSeq | - | 0.90 | 45 | 22 | 13 | 18 | 52 | 12 | 0 | 0 |
| NC_034325 | MiSeq | - | 0.91 | 51 | 17 | 13 | 13 | 44 | 11 | 0 | 3 |
| NC_034326 | MiSeq | - | 0.86 | 62 | 23 | 10 | 18 | 50 | 16 | 0 | 0 |
| NC_028719 | HiSeq 2000 | - | 0.93 | 57 | 7 | 8 | 9 | 39 | 25 | 0 | 0 |
| NC_024643 | HiSeq 2500 | CLC Asm. | 0.93 | 47 | 9 | 9 | 8 | 28 | 31 | 0 | 0 |
| NC_017602 | HiSeq 2000 | - | 0.03 | 0 | 0 | 0 | 0 | 0 | 0 | 0 | 0 |
| NC_029943 | NextSeq 500 | CLC Asm. | 0.87 | 60 | 16 | 14 | 16 | 56 | 21 | 0 | 0 |
| NC_029413 | HiSeq 2000 | Geneious | 0.87 | 62 | 8 | 11 | 9 | 31 | 40 | 0 | 0 |
| NC_030212 | HiSeq 2000 | - | 0.92 | 44 | 17 | 13 | 12 | 47 | 16 | 0 | 2 |
| NC_033910 | HiSeq 2000 | PriceTI | 0.89 | 67 | 23 | 14 | 22 | 51 | 11 | 0 | 0 |
| NC_027161 | HiSeq 2500 | CLC Asm. | 0.94 | 53 | 28 | 7 | 26 | 49 | 8 | 0 | 2 |
| NC_035237 | MiSeq | Velvet | 0.86 | 44 | 13 | 12 | 10 | 45 | 33 | 0 | 0 |
| NC_028185 | GA | SPAdes | 0.73 | 21 | 2 | 12 | 2 | 34 | 14 | 2 | 1 |
| NC_019601 | GA IIx | - | 0.90 | 46 | 17 | 9 | 17 | 53 | 19 | 0 | 0 |
| NC_018766 | HiSeq 2000 | YASRA | 0.76 | - | - | - | - | 6 | 0 | 206 | - |
| NC_019602 | MiSeq | - | 0.95 | 52 | 20 | 12 | 10 | 62 | 8 | 1 | 0 |
| NC_022868 | HiSeq 2000 | CLC Asm. | 0.95 | 40 | 9 | 10 | 6 | 49 | 21 | 0 | 0 |
| NC_024038 | NextSeq 500 | Velvet | 0.84 | - | - | - | - | 41 | 31 | 1 | - |
| NC_026301 | HiSeq 2000 | Velvet | 0.95 | 34 | 14 | 5 | 11 | 28 | 9 | 0 | 0 |
| NC_023215 | MiSeq | - | 0.93 | 43 | 20 | 11 | 23 | 80 | 20 | 0 | 0 |
| NC_029819 | MiSeq | YASRA | 0.89 | - | - | - | - | 45 | 14 | 130 | - |
| NC_007977 | GA II | - | 0.88 | 50 | 29 | 13 | 27 | 33 | 20 | 0 | 0 |
| NC_030275 | HiSeq 2000 | SOAPdenovo | 0.96 | 42 | 21 | 15 | 21 | 41 | 15 | 0 | 2 |
| NC_030173 | HiSeq 2000 | SOAPdenovo | 0.95 | 41 | 19 | 10 | 13 | 47 | 13 | 0 | 0 |
| NC_026909 | HiSeq 2000 | GS denovo Asm. | 0.94 | 61 | 27 | 12 | 24 | 47 | 67 | 0 | 1 |
| NC_024066 | HiSeq 2000 | - | 0.95 | 47 | 15 | 16 | 12 | 45 | 16 | 0 | 0 |
| NC_024060 | HiSeq 2000 | - | 0.93 | 59 | 22 | 12 | 22 | 62 | 15 | 0 | 0 |
| NC_028032 | NextSeq 500 | SPAdes | 0.91 | 55 | 18 | 12 | 19 | 45 | 23 | 0 | 0 |

Continued on next page

Supplementary Table 2 – continued from previous page

| Sample | Metadata | | E-score | WRSD | | | | | | Asm. quality | |
|---|---|---|---|---|---|---|---|---|---|---|---|
| | Seq. platform | Asm. software | | LSC | $IR_B$ | SSC | $IR_A$ | Cod. | Non. | Ns | Mism. |
| NC_034702 | MiSeq | - | 0.94 | 53 | 14 | 11 | 8 | 66 | 17 | 0 | 0 |
| NC_034703 | MiSeq | - | 0.95 | 62 | 8 | 14 | 7 | 52 | 16 | 0 | 0 |
| NC_034685 | MiSeq | - | 0.94 | 64 | 7 | 10 | 1 | 50 | 11 | 2 | 0 |
| NC_034699 | MiSeq | - | 0.90 | 60 | 7 | 14 | 9 | 44 | 15 | 0 | 0 |
| NC_034689 | MiSeq | - | 0.94 | 57 | 5 | 10 | 7 | 51 | 24 | 0 | 0 |
| NC_026703 | HiSeq 2500 | - | 0.96 | 31 | 22 | 3 | 17 | 52 | 10 | 0 | 32 |
| NC_034670 | HiSeq 2500 | - | 0.92 | 47 | 18 | 7 | 18 | 52 | 22 | 0 | 2 |
| NC_034694 | MiSeq | - | 0.94 | 50 | 16 | 9 | 18 | 49 | 10 | 87 | 4 |
| NC_034691 | MiSeq | - | 0.93 | 67 | 14 | 8 | 16 | 56 | 19 | 0 | 0 |
| NC_028617 | HiSeq 2500 | - | 0.97 | 38 | 18 | 8 | 8 | 51 | 9 | 0 | 0 |
| NC_028190 | HiSeq 2000 | - | 0.96 | - | - | - | - | 32 | 12 | 0 | - |
| NC_035068 | HiSeq 2000 | ABySS | 0.95 | - | - | - | - | 49 | 7 | 0 | - |
| NC_007578 | HiSeq X Ten | - | 0.85 | 49 | 6 | 11 | 5 | 34 | 22 | 0 | 0 |
| NC_034700 | MiSeq | - | 0.95 | 57 | 12 | 2 | 2 | 45 | 9 | 0 | 0 |
| NC_024064 | HiSeq 2000 | - | 0.93 | 50 | 21 | 14 | 20 | 54 | 9 | 0 | 0 |
| NC_024065 | HiSeq 2000 | - | 0.92 | 54 | 24 | 12 | 24 | 60 | 10 | 0 | 0 |
| NC_027675 | HiSeq 2000 | CLC Asm. | 0.93 | 58 | 24 | 13 | 21 | 64 | 15 | 0 | 0 |
| NC_035238 | HiSeq 2500 | Velvet | 0.91 | 61 | 13 | 12 | 14 | 42 | 24 | 0 | 0 |
| NC_002694 | NextSeq 500 | - | 0.92 | 50 | 9 | 12 | 13 | 59 | 15 | 0 | 0 |
| NC_030540 | MiSeq | - | 0.94 | 31 | 19 | 6 | 4 | 54 | 11 | 0 | 23 |
| NC_034687 | MiSeq | - | 0.93 | 58 | 19 | 14 | 15 | 69 | 15 | 2 | 0 |
| NC_031163 | HiSeq 2500 | - | 0.90 | 48 | 25 | 14 | 25 | 56 | 18 | 0 | 0 |
| NC_035239 | HiSeq 2500 | Velvet | 0.93 | 61 | 13 | 8 | 10 | 38 | 27 | 0 | 0 |
| NC_023256 | HiSeq 2000 | Velvet | 0.93 | 33 | 20 | 8 | 19 | 48 | 13 | 0 | 0 |
| NC_028720 | GA II | - | 0.95 | 40 | 10 | 6 | 6 | 49 | 19 | 0 | 0 |
| NC_028721 | GA II | - | 0.93 | 57 | 17 | 10 | 14 | 60 | 31 | 0 | 2 |
| NC_034698 | MiSeq | - | 0.96 | - | - | - | - | 44 | 6 | 0 | - |
| NC_032724 | HiSeq 2500 | - | 0.94 | 47 | 8 | 13 | 9 | 29 | 23 | 0 | 0 |
| NC_035143 | HiSeq 2500 | Velvet | 0.97 | 36 | 24 | 10 | 20 | 52 | 8 | 0 | 0 |
| NC_015401 | MiSeq | - | 0.95 | 43 | 20 | 3 | 15 | 40 | 7 | 0 | 0 |
| NC_027460 | HiSeq 2500 | - | 0.93 | 55 | 23 | 9 | 14 | 52 | 21 | 0 | 0 |
| NC_027461 | HiSeq 2500 | - | 0.97 | 36 | 10 | 10 | 4 | 40 | 17 | 0 | 0 |
| NC_034763 | HiSeq 2500 | - | 0.96 | 61 | 8 | 7 | 3 | 47 | 6 | 0 | 0 |
| NC_027462 | HiSeq 2500 | - | 0.97 | 53 | 12 | 9 | 4 | 37 | 13 | 0 | 0 |
| NC_016927 | HiSeq 2500 | - | 0.97 | 47 | 14 | 9 | 5 | 61 | 7 | 0 | 0 |
| NC_034765 | HiSeq 2500 | - | 0.97 | 53 | 6 | 6 | 5 | 49 | 4 | 0 | 0 |
| NC_027463 | GA II | - | 0.83 | 38 | 17 | 3 | 15 | 37 | 14 | 0 | 0 |
| NC_027676 | HiSeq 2000 | CLC Asm. | 0.95 | - | - | - | - | 50 | 8 | 0 | - |
| NC_034764 | HiSeq 2500 | - | 0.96 | 50 | 12 | 6 | 4 | 37 | 7 | 0 | 0 |
| NC_017835 | HiSeq X Ten | - | 0.97 | 51 | 14 | 9 | 6 | 29 | 11 | 0 | 0 |
| NC_031333 | MiSeq | CLC Asm. | 0.96 | 39 | 18 | 9 | 1 | 52 | 9 | 0 | 0 |
| NC_028349 | HiSeq 2500 | - | 0.89 | 55 | 10 | 13 | 14 | 39 | 31 | 0 | 4 |
| NC_030493 | HiSeq 2000 | Velvet | 0.93 | 49 | 18 | 10 | 15 | 52 | 6 | 0 | 0 |
| NC_024067 | HiSeq 2000 | - | 0.92 | 50 | 22 | 6 | 20 | 54 | 16 | 0 | 0 |
| NC_031194 | HiSeq 2000 | Velvet | 0.90 | 52 | 32 | 4 | 32 | 58 | 13 | 0 | 0 |
| NC_030756 | HiSeq 2500 | CLC Asm. | 0.96 | 49 | 20 | 12 | 21 | 59 | 6 | 0 | 0 |
| NC_009259 | HiSeq 4000 | - | 0.95 | 50 | 10 | 9 | 10 | 65 | 20 | 0 | 0 |
| NC_013991 | HiSeq 2500 | - | 0.91 | 45 | 17 | 13 | 6 | 35 | 18 | 0 | 0 |
| NC_015817 | HiSeq X Ten | - | 0.84 | 42 | 9 | 10 | 12 | 21 | 11 | 0 | 0 |
| NC_021456 | HiSeq 2500 | - | 0.85 | - | - | - | - | 57 | 30 | 0 | - |
| NC_032367 | HiSeq 2500 | - | 0.91 | - | - | - | - | 53 | 28 | 0 | - |
| NC_032366 | HiSeq 2500 | - | 0.89 | - | - | - | - | 44 | 17 | 0 | - |
| NC_028594 | HiSeq 2000 | - | 0.87 | - | - | - | - | 40 | 14 | 0 | - |

Continued on next page

Supplementary Table 2 – continued from previous page

| Sample | Metadata | | E-score | WRSD | | | | | | Asm. quality | |
|---|---|---|---|---|---|---|---|---|---|---|---|
| | Seq. platform | Asm. software | | LSC | $IR_B$ | SSC | $IR_A$ | Cod. | Non. | Ns | Mism. |
| NC_034684 | MiSeq | - | 0.92 | 54 | 13 | 13 | 11 | 61 | 21 | 0 | 0 |
| NC_021439 | HiSeq X Ten | - | 0.96 | - | - | - | - | 23 | 9 | 0 | - |
| NC_035069 | HiSeq 2000 | ABySS | 0.95 | - | - | - | - | 36 | 6 | 0 | - |
| NC_021440 | MiSeq | - | 0.91 | - | - | - | - | 59 | 30 | 0 | - |
| NC_034697 | MiSeq | - | 0.93 | 54 | 19 | 10 | 14 | 64 | 20 | 2 | 0 |
| NC_034692 | MiSeq | - | 0.95 | 55 | 22 | 12 | 9 | 50 | 15 | 0 | 0 |
| NC_034998 | HiSeq 2000 | - | 0.97 | 36 | 11 | 12 | 7 | 27 | 21 | 0 | 0 |
| NC_027276 | MiSeq | - | 0.50 | 0 | 0 | 0 | 0 | 0 | 0 | 91 | 0 |
| NC_008235 | HiSeq 2000 | - | 0.94 | 54 | 29 | 9 | 28 | 58 | 8 | 0 | 0 |
| NC_027425 | HiSeq 2500 | CLC Asm. | 0.19 | 0 | 0 | 0 | 0 | 0 | 0 | 0 | 0 |
| NC_028504 | HiSeq 2000 | CLC Asm. | 0.96 | 49 | 17 | 8 | 9 | 47 | 17 | 0 | 0 |
| NC_009143 | HiSeq 2500 | - | 0.96 | 50 | 31 | 9 | 17 | 53 | 11 | 0 | 0 |
| NC_026291 | HiSeq 2500 | - | 0.95 | 31 | 11 | 5 | 11 | 52 | 19 | 0 | 0 |
| NC_031428 | HiSeq 2000 | - | 0.84 | 52 | 14 | 8 | 17 | 32 | 34 | 0 | 55 |
| NC_034696 | MiSeq | - | 0.95 | 53 | 18 | 15 | 13 | 58 | 15 | 2 | 0 |
| NC_023956 | HiSeq 2000 | SOAPdenovo | 0.90 | 51 | 25 | 11 | 26 | 45 | 27 | 0 | 0 |
| NC_023798 | HiSeq 4000 | SOAPdenovo | 0.90 | 54 | 12 | 9 | 6 | 41 | 37 | 0 | 0 |
| NC_026980 | MiSeq | Geneious | 0.97 | 42 | 15 | 12 | 4 | 49 | 21 | 0 | 0 |
| NC_035240 | HiSeq 2500 | Velvet | 0.92 | 65 | 9 | 13 | 10 | 37 | 21 | 0 | 0 |
| NC_027262 | HiSeq 2500 | - | 0.95 | 36 | 14 | 10 | 19 | 50 | 19 | 0 | 1 |
| NC_034951 | HiSeq 2500 | CLC Asm. | 0.94 | 58 | 28 | 11 | 25 | 48 | 11 | 0 | 0 |
| NC_028069 | HiSeq 2500 | - | 0.95 | 57 | 27 | 14 | 22 | 50 | 10 | 0 | 0 |
| NC_030207 | HiSeq 2000 | - | 0.78 | 47 | 9 | 11 | 7 | 47 | 20 | 1 | 6 |
| NC_023800 | HiSeq 2000 | SOAPdenovo | 0.94 | 38 | 13 | 2 | 11 | 54 | 15 | 1 | 1 |
| NC_029416 | HiSeq 2000 | Velvet | 0.94 | 49 | 15 | 7 | 14 | 44 | 18 | 0 | 0 |
| NC_029825 | MiSeq | YASRA | 0.90 | - | - | - | - | 51 | 20 | 551 | - |
| NC_029822 | MiSeq | YASRA | 0.97 | - | - | - | - | 35 | 7 | 0 | - |
| NC_029823 | MiSeq | YASRA | 0.91 | - | - | - | - | 47 | 15 | 102 | - |
| NC_029824 | MiSeq | YASRA | 0.93 | - | - | - | - | 42 | 7 | 56 | - |
| NC_032057 | MiSeq | Velvet | 0.86 | 52 | 16 | 16 | 10 | 38 | 28 | 0 | 2 |
| NC_035066 | HiSeq 2000 | ABySS | 0.95 | - | - | - | - | 32 | 17 | 0 | - |
| NC_020098 | HiSeq 4000 | - | 0.95 | 46 | 19 | 9 | 15 | 52 | 9 | 0 | 0 |
| NC_034362 | MiSeq | - | 0.94 | 53 | 7 | 9 | 4 | 45 | 5 | 0 | 2 |
| NC_019616 | GA IIx | GS denovo Asm. | 0.69 | 34 | 0 | 1 | 0 | 0 | 1 | 0 | 0 |
| NC_027155 | HiSeq 1000 | - | 0.88 | - | - | - | - | 35 | 8 | 0 | - |
| NC_018051 | HiSeq 4000 | MIRA | 0.90 | 46 | 17 | 7 | 10 | 39 | 27 | 0 | 31 |
| NC_031383 | HiSeq 4000 | - | 0.96 | 55 | 29 | 12 | 22 | 46 | 15 | 0 | 0 |
| NC_023790 | HiSeq 2500 | SOAPdenovo | 0.95 | 53 | 23 | 11 | 16 | 48 | 15 | 0 | 0 |
| NC_027677 | HiSeq 2000 | - | 0.92 | - | - | - | - | 41 | 9 | 0 | - |

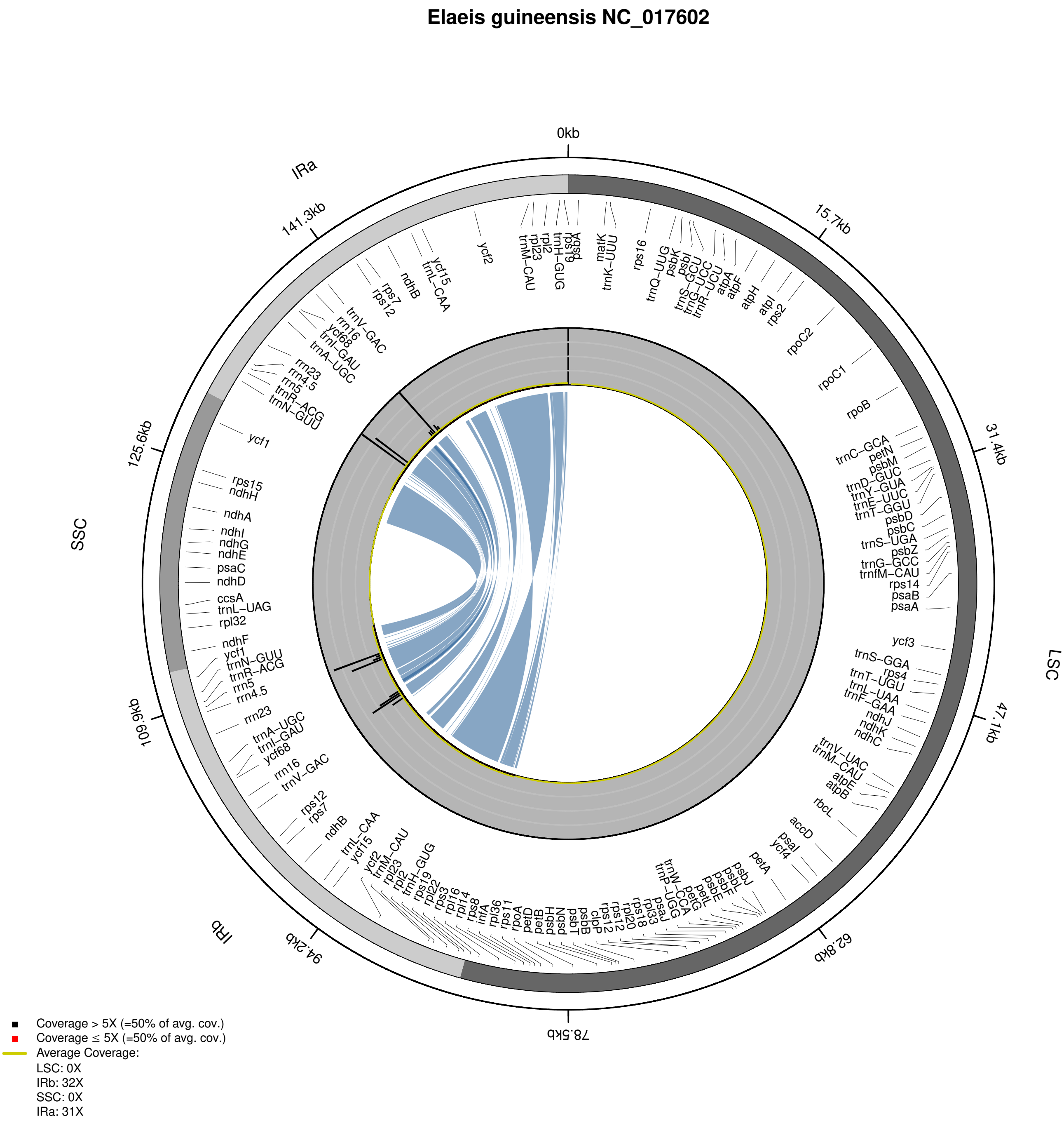


**Supplementary Figure 1:** Distribution of coverage depth in the plastid genome of *Elaeis guineensis* (NC_017602).

**Dioscorea cayenensis subsp. rotundata NC_024170**

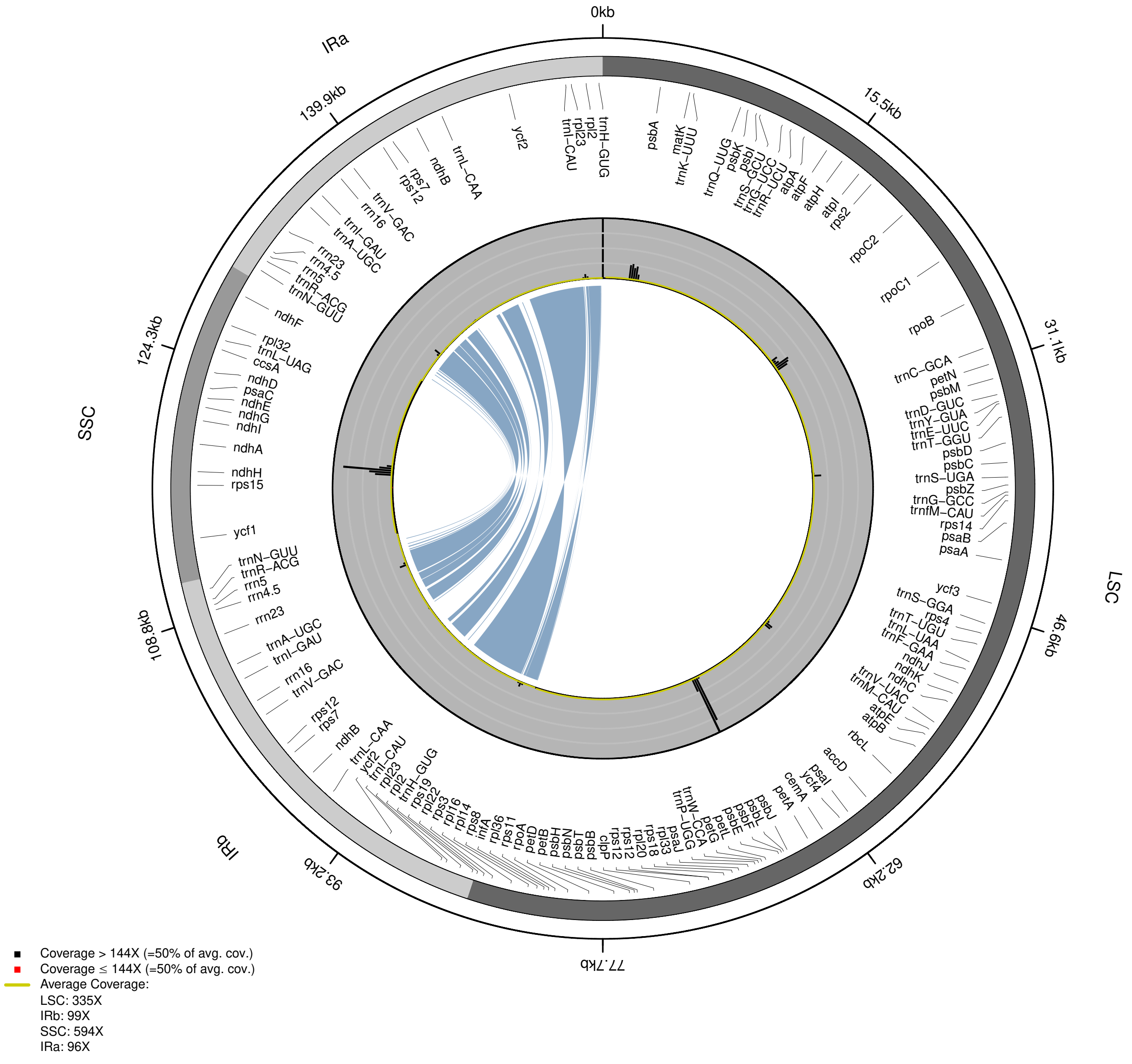


**Supplementary Figure 2:** Distribution of coverage depth in the plastid genome of *Dioscorea cayenensis* subsp. *rotundata* (NC_024170).

**Populus tremula NC_027425**

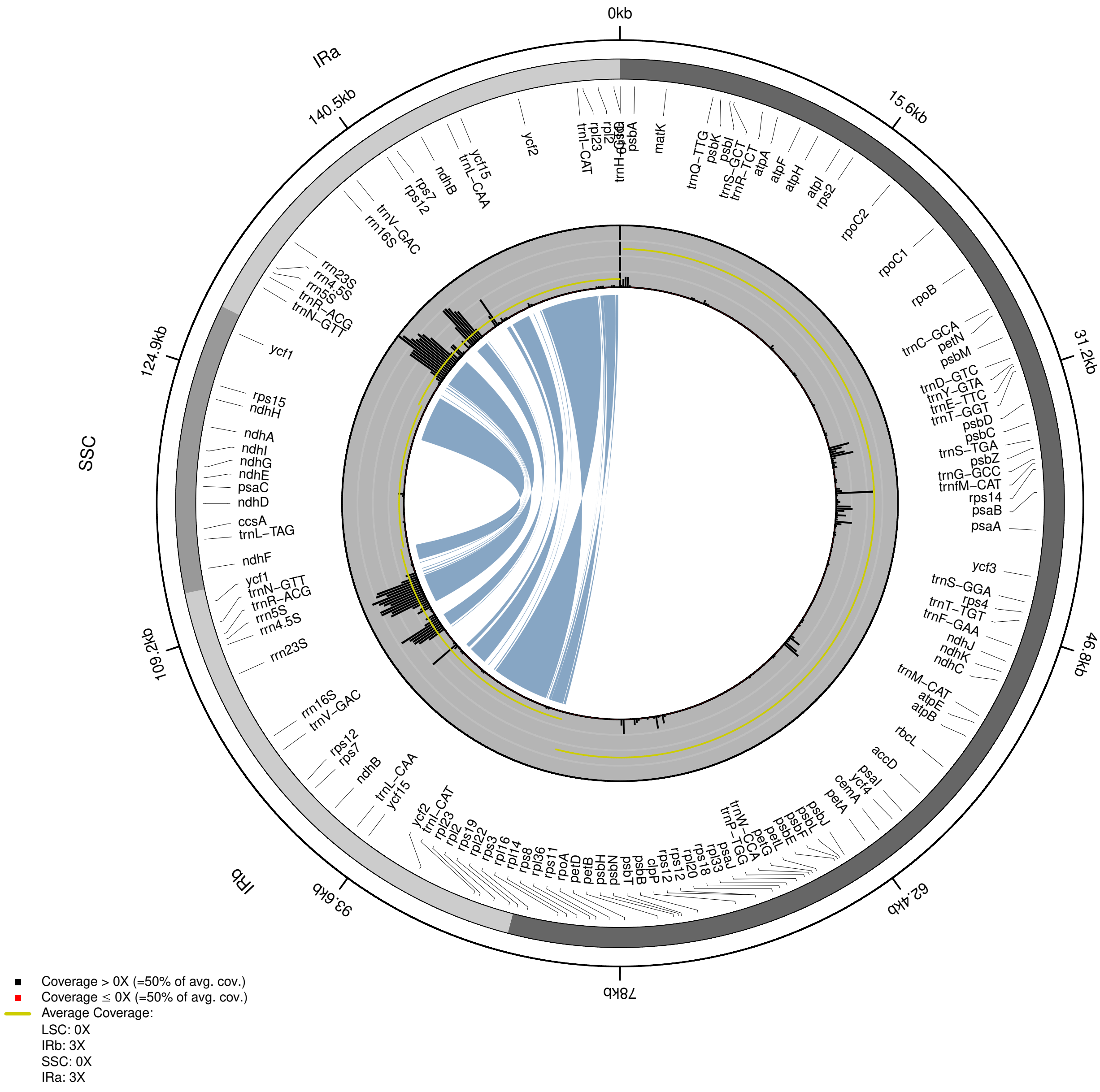


**Supplementary Figure 3:** Distribution of coverage depth in the plastid genome of *Populus tremula* (NC_027425).

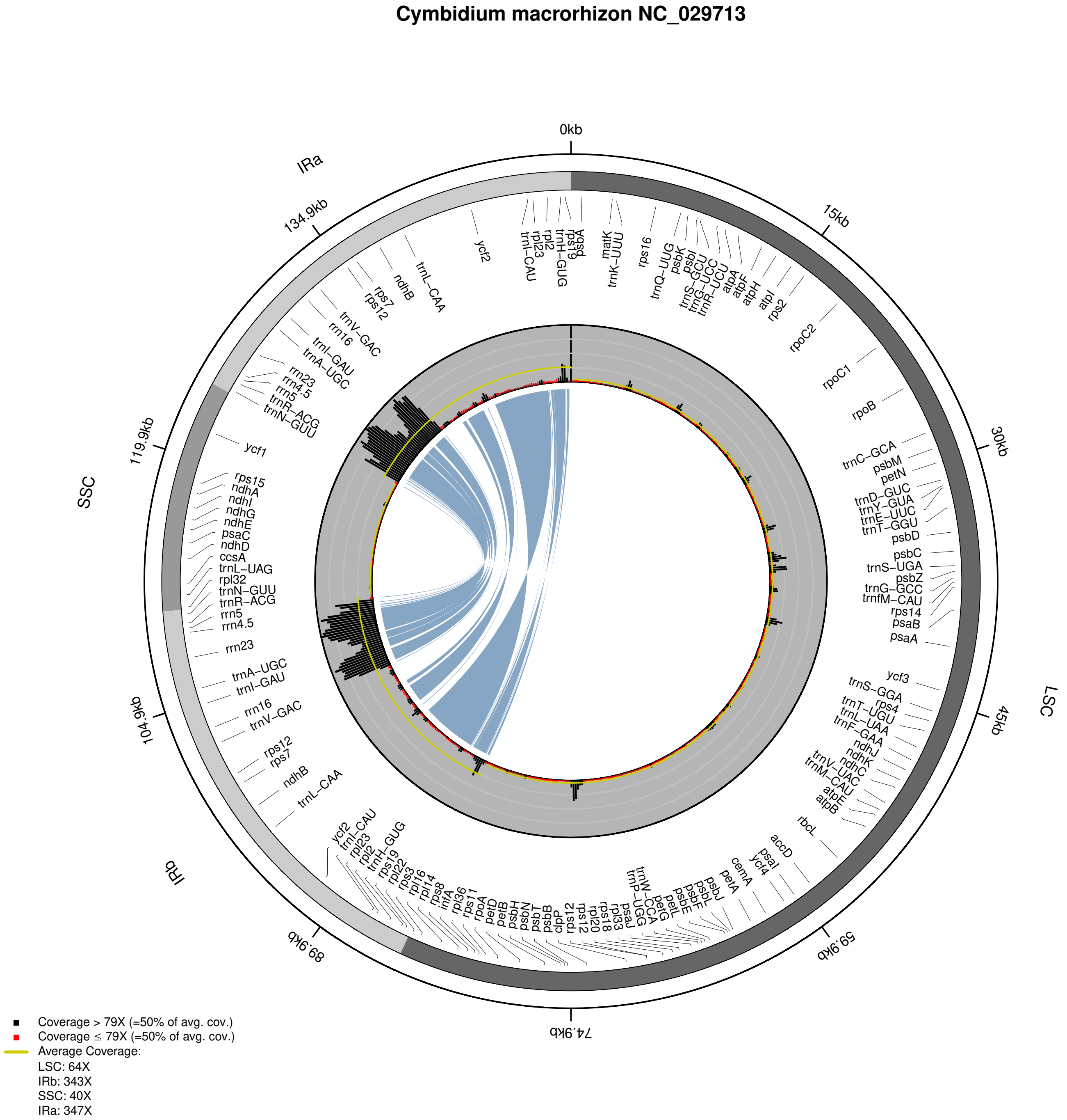


**Supplementary Figure 4:** Distribution of coverage depth in the plastid genome of *Cymbidium macrorhizon* (NC_029713).

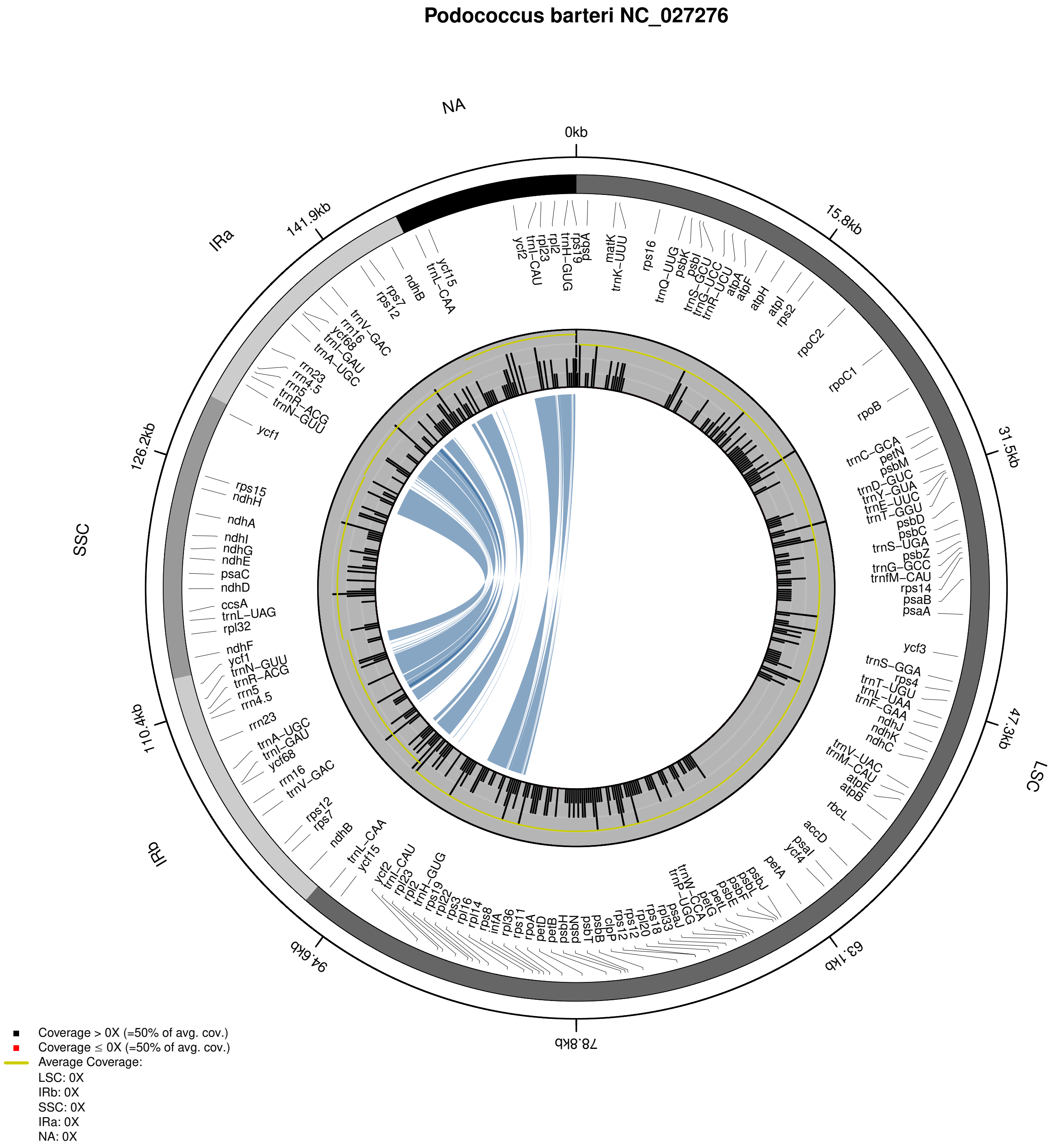


**Supplementary Figure 5:** Distribution of coverage depth in the plastid genome of *Podococcus barteri* (NC_027276).

**Cannabis sativa NC_026562**

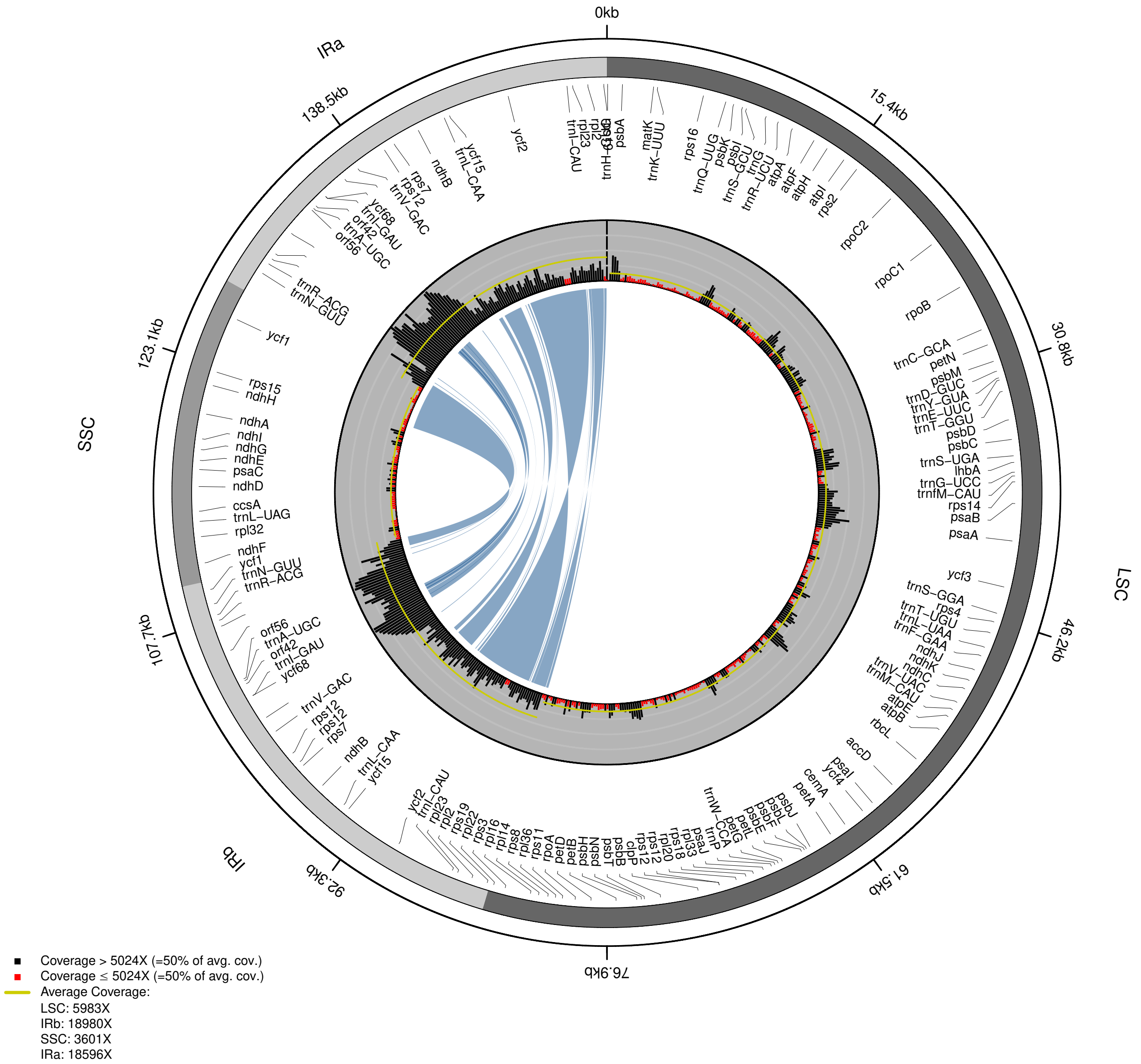


**Supplementary Figure 6:** Distribution of coverage depth in the plastid genome of *Cannabis sativa* (NC_026562).

**Vaccinium macrocarpon NC_019616**

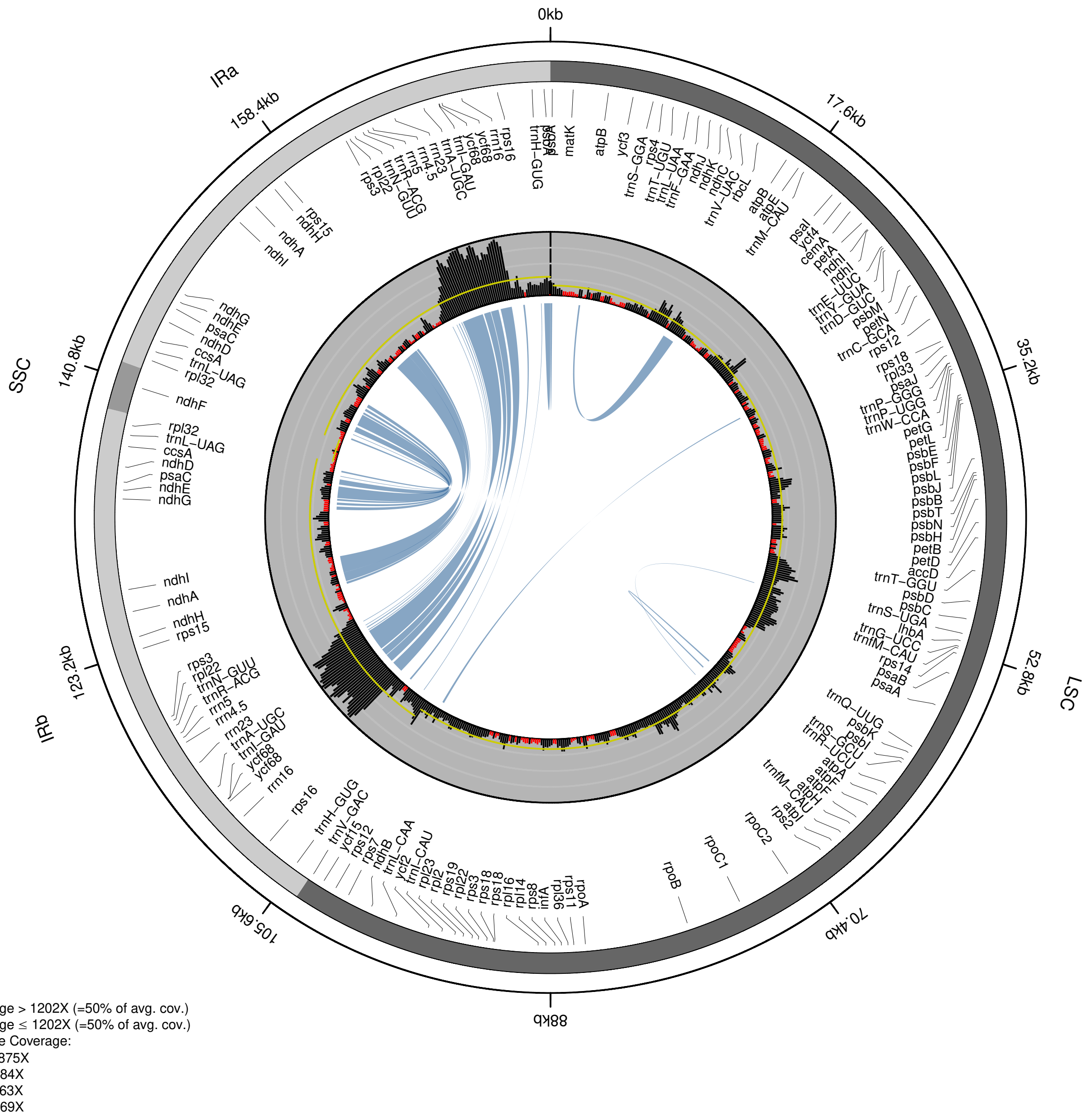


**Supplementary Figure 7:** Distribution of coverage depth in the plastid genome of *Vaccinium macrocarpon* (NC_019616).

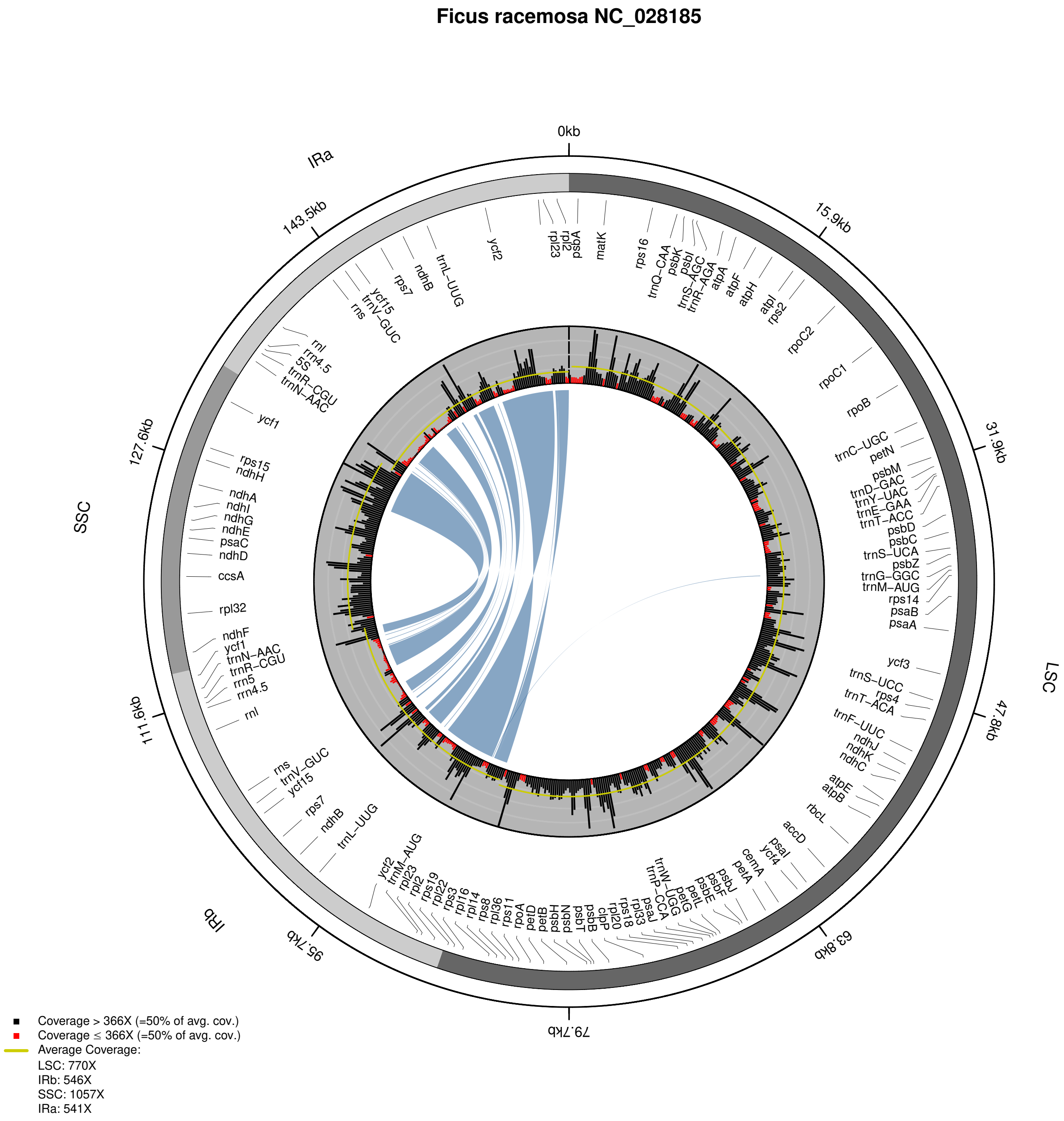


**Supplementary Figure 8:** Distribution of coverage depth in the plastid genome of *Ficus racemosa* (NC_028185).

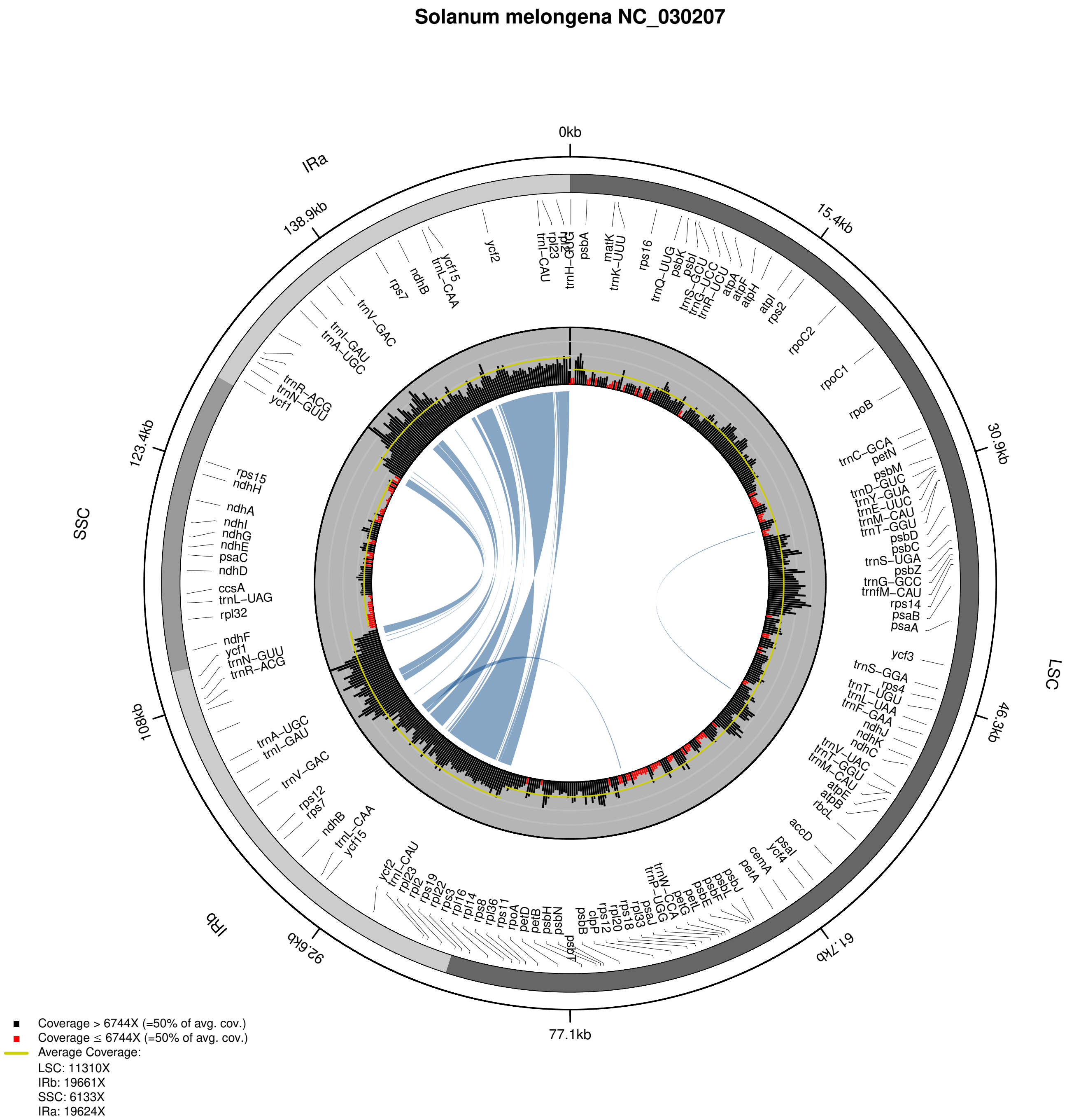


**Supplementary Figure 9:** Distribution of coverage depth in the plastid genome of *Solanum melongena* (NC_030207).

**Brassica napus NC_016734**

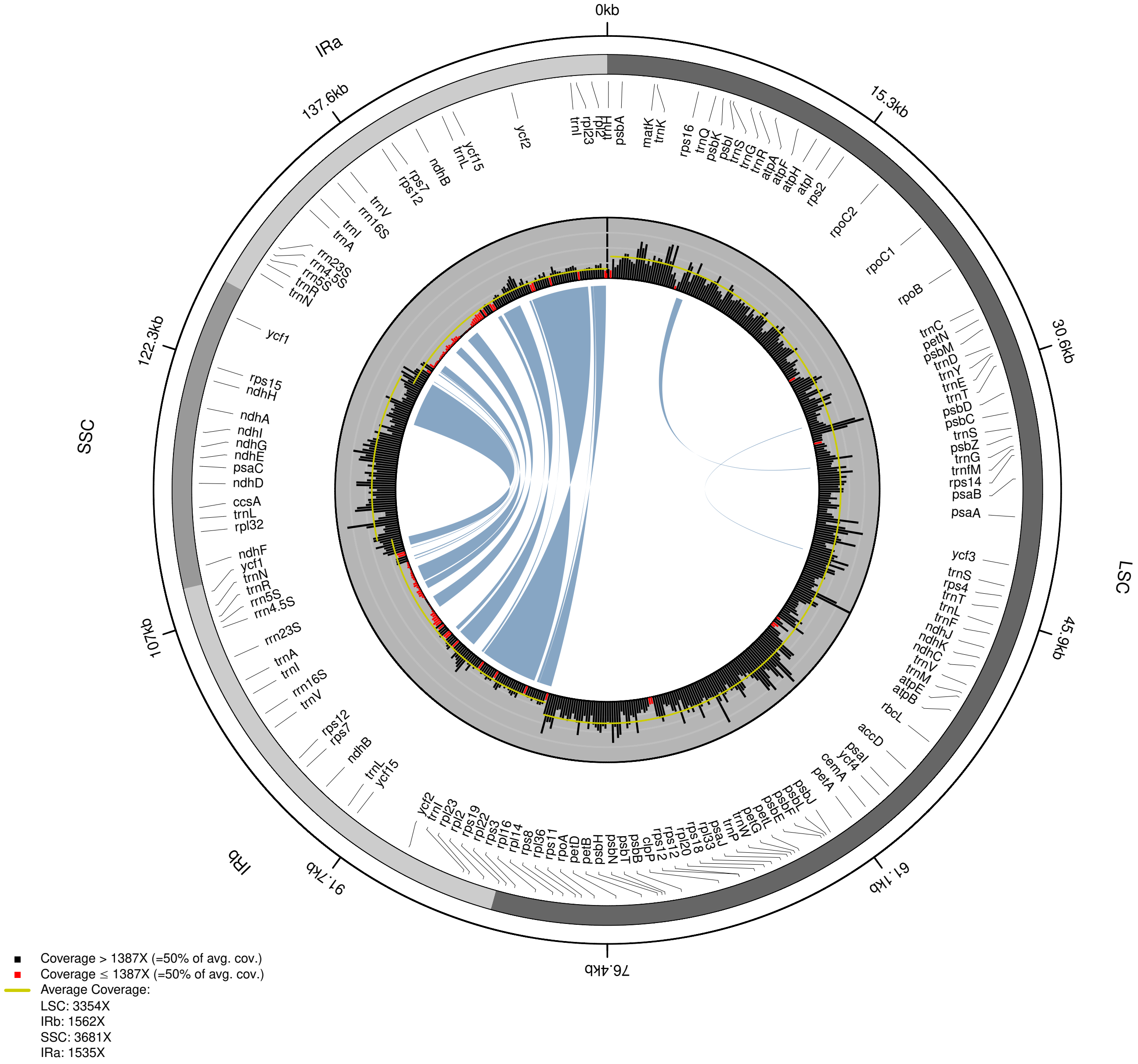


**Supplementary Figure 10:** Distribution of coverage depth in the plastid genome of *Brassica napus* (NC_016734).

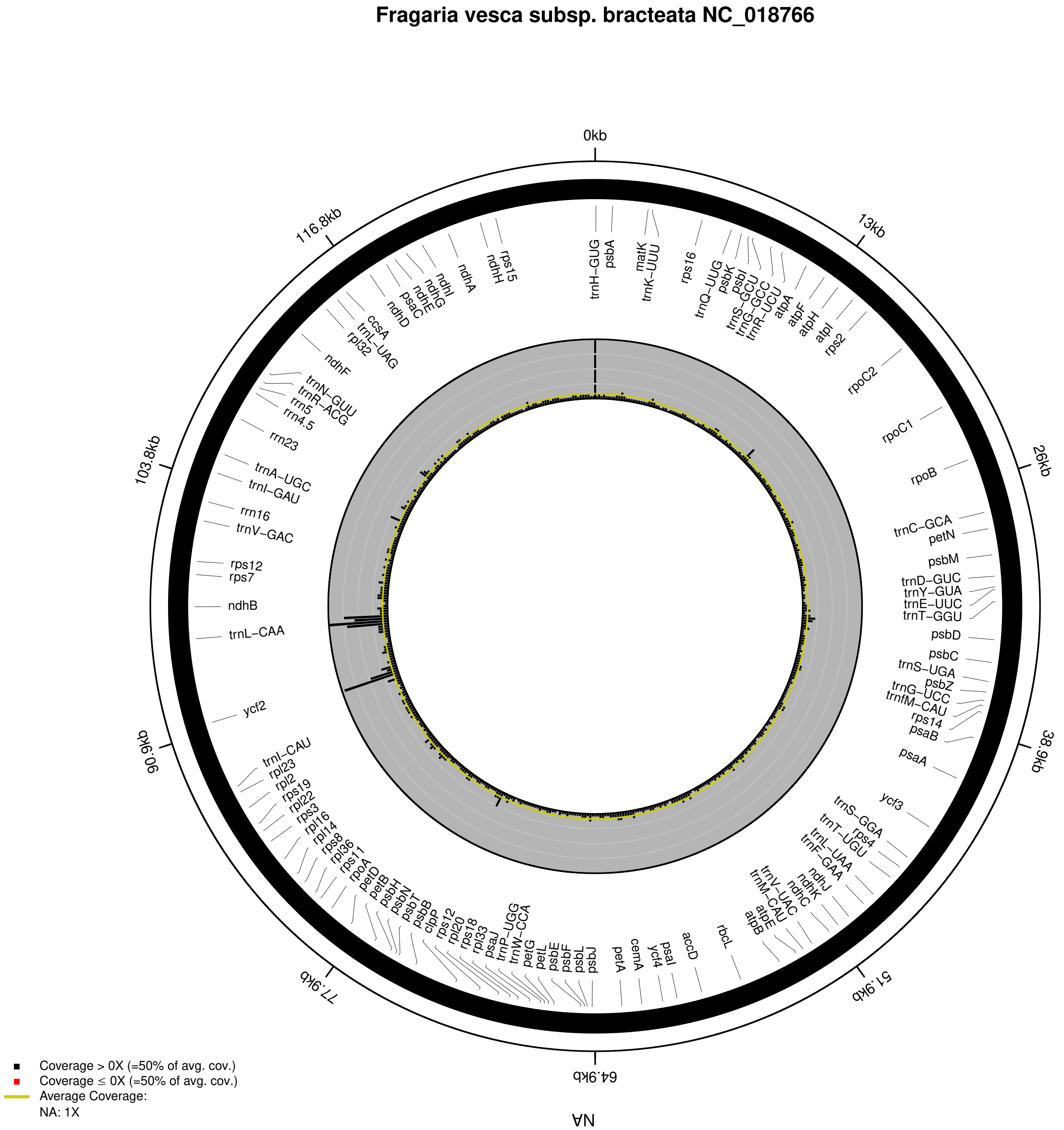


**Supplementary Figure 11:** Distribution of coverage depth in the plastid genome of *Fragaria vesca* subsp. *bracteata* (NC_018766).

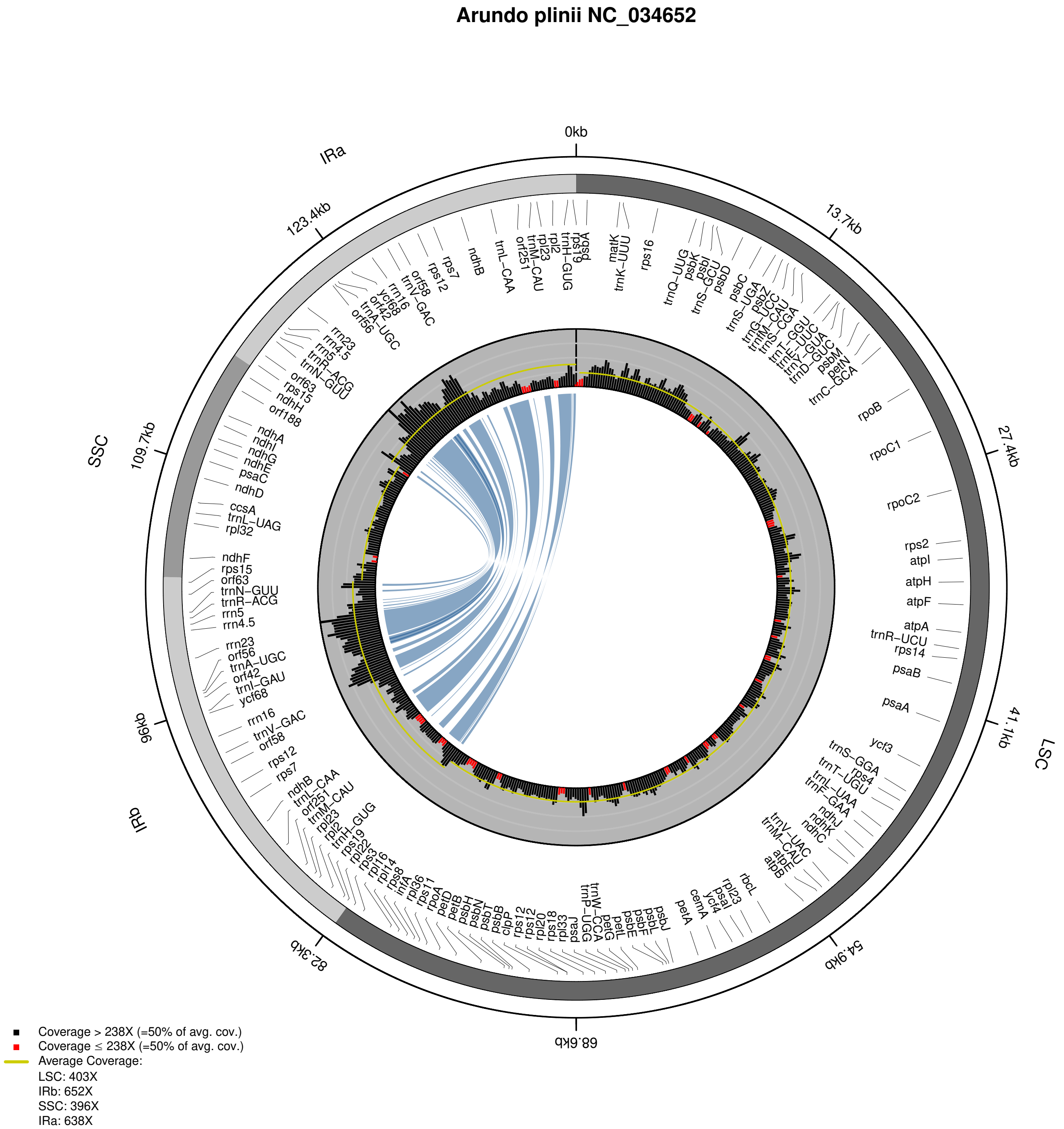


**Supplementary Figure 12:** Distribution of coverage depth in the plastid genome of *Arundo plinii* (NC_034652).

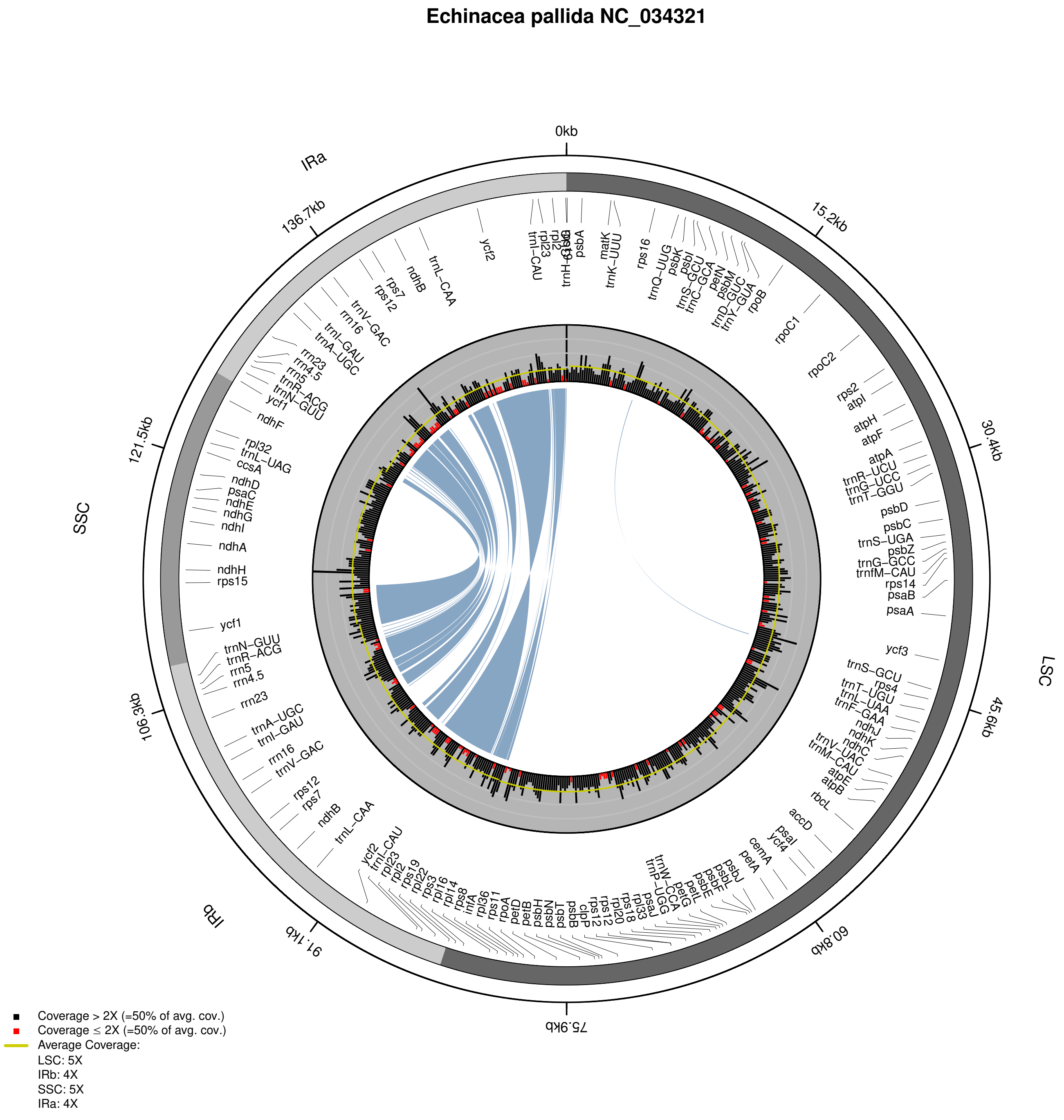


**Supplementary Figure 13:** Distribution of coverage depth in the plastid genome of *Echinacea pallida* (NC_034321).
